# Native trees within plantations and surrounding forest cover are essential for bird conservation in cashew-dominated landscapes within a biodiversity hotspot

**DOI:** 10.1101/2024.11.30.625592

**Authors:** Nandita Madhu, Vishal Sadekar, Nayantara Biswas, Rajah Jayapal, Anushka Rege, Rohit Naniwadekar

## Abstract

1. Annually, agriculture is responsible for about 60% of tropical forest loss globally.
2. strategies in agricultural landscapes range from enhancing farm-level habitat diversity to retaining forest cover within landscapes. A critical question is determining the relative influence of native trees within farms and forest cover around them on avian diversity, trait responses and species persistence in tree-based systems in landscapes where protected areas are often absent, often the case in lowland tropical forest ecosystems.
3. By sampling birds across gradients of cashew plantation intensification and varying levels of surrounding cashew and forest cover, we examined the relative influence of site- and landscape-level factors on species composition, taxonomic, functional, and phylogenetic diversity, as well as species- and trait-specific responses, and whether phylogenetic relationships among bird species influenced these responses.
4. At the community level, forests, cashew agroforests, and cashew monocultures hosted compositionally different bird communities. Site-level variables, specifically the proportion of native trees, influenced taxonomic and phylogenetic diversity, while landscape-level variables had no effect. None of the predictor variables accounted for the variation in functional diversity.
5. At the species level, site-level variables, specifically the proportion of native trees, and landscape-level variables, particularly cashew cover around plantations, had positive and negative impacts on species responses, respectively. At the trait level, evergreen forest specialists were associated positively with the proportion of native trees and negatively associated with cashew cover. Resident species had a positive association with forest cover. We found a phylogenetic signal in species responses to predictor variables, indicating that closely-related species responded similarly to predictor variables.
6. *Synthesis and applications*: To retain taxonomic and phylogenetic diversity and ensure persistence of evergreen forest specialists, it is essential to retain native trees within cashew plantations and maintain surrounding forest cover. Future policies should incentivise the retention and restoration of forest patches and restrict the felling of native trees within cashew plantations.

## INTRODUCTION

Annually up to 8.8 Mha of tropical forests were deforested globally for agriculture from 2011 to 2015 (Pendrill et al., 2022). Rapid expansion of monocultures and agroforestry plantations are significant drivers of tropical forest loss (Jaureguiberry et al., 2022). Despite the role of protected areas in biodiversity conservation, they are insufficient for tropical biodiversity conservation due to their limited coverage (Baldwin and Beazley, 2019), isolation, and skewed distribution in higher elevations (Elsen et al., 2018). It is imperative to make human- dominated landscapes more biodiversity-friendly. At the landscape level, retaining forest patches in agroforestry-dominated landscapes provides breeding and foraging habitats for animals (Muñoz-Sáez et al., 2020; Ranganathan et al., 2010). At the site level, retaining native trees within agroforests can offer favourable microclimates and resources in modified landscapes (Raman et al., 2021). However, the relative influence of landscape- and site-level drivers on biodiversity is poorly understood. Existing information is primarily from temperate regions and vineyards, with lower representation from tropical agroforests (Birch et al., 2024). This information is critical for informing biodiversity-friendly practices in plantation- dominated landscapes in tropics.

Relationships between taxonomic, functional and phylogenetic diversity with predictor variables may differ. Moreover, drivers of diversity may differ from drivers of threatened species occurrence, both important for conservation objectives. As aggregated metrics, community measures can obscure nuanced responses of species or functional groups to environmental conditions (Neilly et al., 2016). This makes it critical to determine drivers at community, trait and species levels.

Taxonomic, functional and phylogenetic diversity measures provide complementary information. Functional and phylogenetic metrics provide information on community stability, resilience, and ecosystem functioning. Properties like dispersion and clustering offer insights into community assembly processes (Bouvier et al., 2024). While studies increasingly examine the impacts of habitat conversion on diversity measures (Rurangwa et al., 2022), research incorporating trait- and species-specific responses alongside them remains limited (but see Aldabe et al., 2023).

Birds are a suitable taxa for studying biodiversity impacts due to their well-resolved taxonomy and phylogeny. Several bird traits are closely linked to their niches (Pigot et al., 2020). Additionally, birds provide critical ecosystem services, such as pollination and seed dispersal (Sekercioglu, 2012, 2006).

Cashew (*Anacardium ocidentale*), a crop from Brazil, was introduced in other countries by the Portuguese in the 16^th^ century (Eapen et al., 2003). Today, India is among the leading cashew nut exporters (Rege and Lee, 2022). The cashew-growing region of Maharashtra state is part of the Western Ghats-Sri Lanka Biodiversity Hotspot that has experienced a 15% increase in cashew cultivation area in the past decade (Chhaya et al. in prep.). A third of two sub-regions of Sindhudurg district is now under cashew cultivation (Rege and Lee, 2022).

Traditionally, cashew was grown among native trees as agroforests. However, these are being replaced by cashew monocultures. Southern Maharashtra has no Protected Areas, but fragmented patches of government- and privately-owned forests still exist that support breeding populations of threatened species, including hornbills, many of which are endemic to the Western Ghats Biodiversity Hotspot (Punjabi and Rao, 2017). The mosaic of forest patches, cashew agroforests, and monocultures at varying distances from forests provides an opportunity to investigate the influence of site- and landscape-level characteristics on bird diversity. Given cashew’s expansion in other biodiversity hotspots (e.g., Guinean Forests of West Africa), understanding the drivers of bird diversity in cashew plantations has global relevance.

This study assessed the relative influence of site- and landscape-level drivers on bird diversity and persistence in a cashew-dominated landscape in India’s Western Ghats biodiversity hotspot. Specifically, we examined 1) differences in bird species composition among forests, cashew agroforests, and cashew monocultures, 2) the relative influence of site- and landscape-level drivers on bird taxonomic, functional, and phylogenetic diversity, 3) the relative influence of site- and landscape-level drivers on species- and trait-specific responses, and 4) a phylogenetic signal in species responses to predictor variables. This is one of the few studies to examine multifaceted responses (species-, trait- and community-level) of birds to local and landscape-level drivers with findings of relevance for landscape and plantation management for biodiversity.

## METHODS

### Study Area

From January to May 2024, we conducted the study in the low-elevation forests and cashew plantations of Dodamarg taluk of Sindhudurg district of Maharashtra state in India (Fig. 1). This is the northern part of the Western Ghats-Sri Lanka Biodiversity Hotspot, known for high levels of species endemism. The average annual rainfall of this region is about 3,500 mm, and the annual temperature ranges between 12–40°C (Munje and Kumar, 2022). The dominant vegetation type in these lower elevations is moist deciduous forest with patches of evergreen forests. The forests harbour trees like *Beilschmiedia dalzellii*, *Knema attenuata*, *Saraca asoca*, *Terminalia elliptica*, *Terminalia paniculata*, and *Careya arborea,* among others (Biswas et al., 2024). Some portions of Reserved Forests are declared Conservation Reserves (Punjabi and Rao, 2017). These forests are critical in harbouring bird diversity (Biswas et al., 2023). Protected Areas closest to the region (Sahyadri Tiger Reserve, Radhanagari Wildlife Sanctuary) are in high elevations.

**Figure 1.**
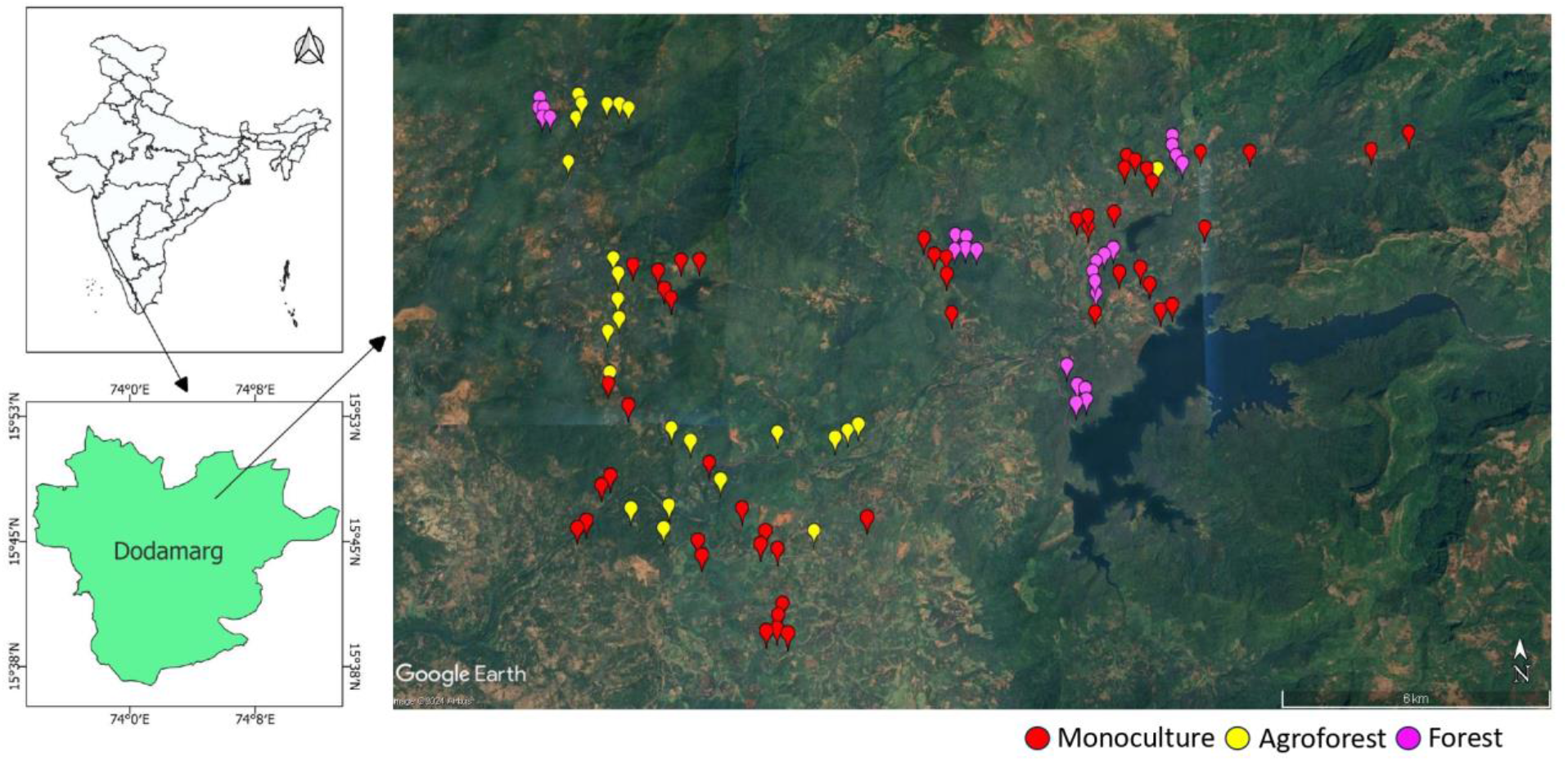
The satellite image displays 100 sampling points (monocultures, agroforests, and forests) in Dodamarg Taluka, India.

### Field Methods

#### Bird sampling

To determine the diversity and occurrence of birds, NM and VS conducted point count surveys at 100 locations. Points were distributed across cashew monocultures (50), cashew agroforests (25) and forests (25) (Fig. 1), separated by at least 250 m to minimise spatial independence (Lee et al., 2024). We sampled them five times between January and April 2024. Sampling was carried out in the mornings between 0645–1000 hr. We recorded species identity and number of individuals seen or heard (except in flyovers), including only those detected within 100 m of the point for analysis.

#### Vegetation sampling

At each point, we established a 10-m radius circular plot, where we recorded the number of woody plants (≥ 30 cm girth at breast height), species identity, girth, tree height, and canopy cover. Using this information, we estimated tree density (ha^-1^), basal area (m^2^ ha^-1^), tree height (m), canopy cover (%), and tree species richness across the three land-use categories (forest, agroforest, and monoculture).

#### Functional trait data

Morphological trait and migratory status (resident/migratory) data were obtained from AVONET (Tobias et al., 2022), dietary trait data from Elton Traits (Wilman et al., 2014), and habitat preferences (evergreen/deciduous) from Ali and Ripley (1999). Beak width, tarsus length and hand wing index are associated with diet and locomotion in birds (Pigot et al., 2020). Similarly, insectivores, frugivores and forest-specialists are negatively impacted by habitat modification (Shahabuddin et al., 2021).

#### Phylogenetic data

We obtained 100 Bayesian phylogenetic trees from birdtree.org (Jetz et al., 2012), following the Ericson backbone {Ericson All Species: a set of 10,000 trees with 9993 Operational Taxonomic Units (OTUs) each}. Using these trees, we obtained a maximum clade credibility tree for further analysis, i.e. phylogenetic diversity estimation and joint species distribution modelling.

#### Landuse landcover data

Apart from site-level factors, the amount and type of habitat at landscape levels impacts bird prevalence (Fahrig, 2013). Using an existing land-use land-cover raster from a recent study, we obtained the amounts of forest and cashew cover around each sampling point within a pre-defined buffer radius (Rege et al., 2022).

#### Analysis

We conducted all the analyses in R version 4.4.0 (R Core Team, 2024).

#### Tree species richness and vegetation structure attributes

To determine differences in tree species richness across land-use categories, we used the Hill Shannon diversity measure (q=1), employing the sample-coverage-based-rarefaction method, using the ‘estimateD’ function of the ‘iNEXT’ package (Hsieh et al., 2015). Roswell et al., (2021) recommend the Hill-Shannon diversity measure that focuses on the diversity of common species to compare diversity across multiple categories. We bootstrapped the data 1000 times to estimate 95% confidence intervals (CIs). We inferred statistical differences between categories if the 95% CI did not overlap (Cumming et al., 2007). To determine differences between site-level vegetation structural attributes like tree density, basal area, tree height, and canopy cover across land-use categories, we used general linear models. We tested for normality of each attribute using Shapiro-Wilk’s test and log-transformed non-normal attributes to approximate normality. To determine differences in proportions of native trees and cashew trees per plot, we used beta regression. We estimated marginal means and assessed pairwise contrasts using Tukey’s post-hoc tests. We used *betareg* and *emmeans* functions of R-packages ‘betareg’ and ‘emmeans’ (Zeileis et al., 2004). We estimated bootstrapped 95% CI of all structural attributes using *boot* and *confint* functions of R-packages ‘boot’ and ‘car’ (Canty and Ripley, 1999; Fox et al., 2001).

#### Determining buffer radius for landscape-level predictors

We followed the best scale-of-effect approach to objectively determine the appropriate buffer radii (Bonfim et al., 2024; Fahrig, 2013). We extracted the forest cover percentages around the 100 sampling points at different buffer radii (from 100 to 3000 m at 100 m intervals). We then determined the explained variation (*R^2^*) for the model examining the relationship between taxonomic species richness and the extent of forest cover at each 100 m interval. The radii at which the *R^2^* values peaked were shortlisted. In our study, the relationship between species richness and forest cover peaked at 100 m, 200 m, and 800 m buffer radii (Fig. S2).

We selected 800 m as the radius for estimating cashew and forest cover, as the 100 and 200 m buffers would capture information similar to our vegetation plots at each point.

#### Selecting local- and landscape-level predictors

We checked for correlations between variables using Pearson’s correlation coefficient (Fig. S3). We found strong correlations (*r* > 0.7) among the proportion of native trees (per plot), the proportion of cashew trees (per plot), tree species richness per plot, tree density per hectare, basal area (m^2^ ha^-1^) and tree height (m). Since our study aimed to determine the role of native trees within agroforests, we chose the proportion of native trees (per plot) and canopy cover for further analysis.

We found that the two landscape-level variables, forest cover and cashew cover, were not strongly correlated with each other or with proportion of native trees and canopy cover (*r* < 0.7). Thus, we used two site-level variables (proportion of native trees and canopy cover) and landscape-level variables (forest cover and cashew cover) for further analyses.

#### Bird composition and diversity

We only included terrestrial birds to analyse bird composition and diversity as our study focused on understanding the impacts of land-use change on terrestrial birds.

We examined differences in bird composition across land-use types (forest, agroforest, and monoculture) using non-metric multidimensional scaling (NMDS) with the Bray-Curtis dissimilarity index. We used the function *metaMDS* from the R package ‘vegan’ on bird abundance data pooled across temporal replicates (Oksanen et al., 2001). We used ecological null models with the *oecosimu* function of the ‘vegan’ package (Oksanen et al., 2001) to validate the NMDS ordination fit, using the ‘swap-count’ method following Dexter et al., 2018. We performed an analysis of similarities (ANOSIM) and a permutational multivariate analysis of variance (PERMANOVA) to test for significant differences between land-use types using functions *anosim* and *adonis2* from the R-package ‘vegan’(Oksanen et al., 2001).

The sample coverage, as estimated using individual-based abundance data, was very high (Forest: 0.9317; Agroforest: 0.9341; Monoculture: 0.9435), indicating sampling adequacy for each category. We did the following to determine the influence of site- and landscape-level factors on taxonomic diversity. First, we estimated the sample coverage for each point. Then, we used the minimum value of the sample coverage per point to calculate the Hill-Shannon Diversity value for each point. We regressed those Hill-Shannon values as a function of local and landscape-level predictors. We used the R package ‘iNEXT’ to estimate sample coverage and Hill-Shannon Diversity.

Since functional diversity is influenced by including multiple correlated traits (De Bello et al., 2021), we checked for correlation between bird traits using Pearson’s correlation and used uncorrelated traits for further analysis (Fig. S4). We chose beak width, tarsus length, Kipp’s distance, all diet-related variables, migratory status (resident/migratory), and habitat preference (evergreen/deciduous) as traits for further analysis. These traits capture information on different dimensions, including diet (beak width and diet-related variables), dispersal ability (hand-wing index) and ecology (migratory status and habitat preference), thereby providing a holistic measure of the functional diversity of birds.

To ensure that we estimate functional and phylogenetic diversity using similar methods, we used the Mean Pairwise Distance (MPD) metric. This tree-based metric averages the functional or phylogenetic distances between all possible species pairs to estimate the observed value of functional or phylogenetic diversity for each sampling point. We used the ‘unweighted pair group’ method for constructing a tree based on functional traits. Since functional and phylogenetic diversity metrics can often be strongly influenced by taxonomic species richness, we estimated standardised effect sizes of the MPD values. We used functions *vegdist*, *hclust*, *as.phylo*, and *ses.mpd* from R packages ‘vegan’, ‘ade4’, and ‘picante’ for this analysis (Dray et al., 2002; Kembel et al., 2010; Oksanen et al., 2001).

To determine the effects of local- (proportion of native trees and canopy cover) and landscape-level (forest cover and cashew cover) drivers on taxonomic diversity, SES-fMPD (functional diversity) and SES-pMPD (phylogenetic diversity), we used general linear models (Gaussian error distribution). These diversity metrics, as estimated for each sampling point, were used for the linear model analysis.

#### Species and trait responses

We used the Hierarchical Modeling of Species Communities (HMSC) framework (Ovaskainen and Abrego, 2020), a Joint Species Distribution Modeling (JSDM) approach, to determine 1) species responses to local- and landscape-level predictors, 2) trait responses to local- and landscape-level predictors, and 3) phylogenetic signal in the residual variation i.e., variation beyond that explained by traits. This analysis complements diversity and composition analyses and allows determining responses of specific species and traits to different environmental and anthropogenic variables.

We excluded rare species (32 species that occurred in less than 5% of the points) from the analysis since they provide little information on community assembly processes and pose problems for Markov Chain Monte Carlo (MCMC) convergence. We used the same bird trait data and phylogenetic tree for this analysis as described earlier. We used the R package ‘Hmsc’ to fit the model in a Bayesian framework using prior distributions (Tikhonov et al., 2019). We modelled species and trait responses as a function of local- (proportion of native trees and canopy cover) and landscape-level (forest cover in 800 m and cashew cover in 800 m) predictors. We incorporated the location of each sampling point as a random effect to account for spatial autocorrelation. Using a presence-absence model with a probit link, we modelled occurrence probabilities of 70 species. We sampled posterior distributions with three MCMC chains, thinned by 1000, to achieve 250 samples for each chain. The first 50,000 samples were removed as burn-in.

## RESULTS

### Site-level habitat characteristics

We enumerated 637 trees of 56 species across our 100 plots (Table S1). Tree diversity in forests was 2.3 and 18 times higher than in cashew agroforests and monocultures, respectively (Fig. S1; Table S2). Forests had higher tree density (Adj. *R^2^* = 0.29, *F2, 78* = 17.27, *p* < 0.001), basal area (Adj. *R^2^* = 0.40, *F2, 78* = 28.25, *p* < 0.001), tree height (Adj. *R^2^* = 0.69, *F2, 78* = 93.89, *p* < 0.001), and canopy cover (Adj. *R^2^* = 0.14, *F2, 78* = 7.539, *p* = 0.001) than cashew monocultures, and higher tree height (Adj. *R^2^* = 0.69, *F2, 78* = 93.89, *p* < 0.001), and canopy cover (Adj. *R^2^* = 0.14, *F2, 78* = 7.539, *p* = 0.001) than cashew agroforests (Fig. S1; Table S3 and S4). Forests had a higher proportion of native trees than agroforests (Pseudo *R^2^* = 0.96).

### Bird composition and diversity

We recorded 6,735 individuals of 102 bird species belonging to 38 families (Table S5). Bird species composition across land-use types was statistically dissimilar (*Ranosim* = 0.58, *p* = 0.001; PERMANOVA *R^2^* = 0.25, df = 2, *F* = 16.08, *p* = 0.001; Fig. 2a). The null hypothesis was rejected by a one-sample z-test after 1000 permutations, indicating differences in community structure (*z* = -1.96, *p* < 0.001). Interestingly, the species composition of cashew agroforests was between forests and cashew monocultures, indicating shared species with the other two categories (Fig. 2a).

**Figure 2.**
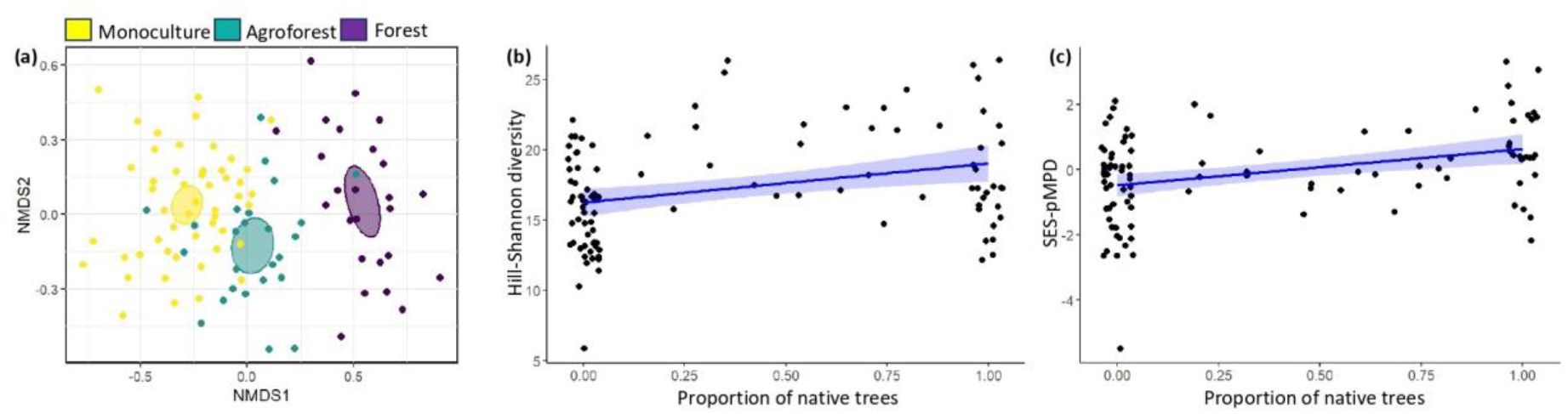
(a) Bird species composition across land-use types, (b) taxonomic diversity (Hill-Shannon diversity) and (c) phylogenetic diversity (SES-pMPD) versus the proportion of native trees in the 10-m circular radius plot around the tree.

We found no association between canopy cover, forest cover, or cashew cover, and taxonomic, functional, or phylogenetic diversity. The proportion of native trees positively influenced taxonomic (Adj. *R^2^* = 0.08, *F4, 95* = 3.29, *p* = 0.01) and phylogenetic (Adj. *R^2^* = 0.11, *F4, 95* = 4.30, *p* < 0.05) diversities (Fig. 2b and c; Table S6 and S7). However, functional diversity was not associated with any predictor (Adj. *R^2^* = -0.02, *F4, 95* = 0.32, *p* > 0.05) (Table S8).

### Species and trait-specific associations with local and landscape predictors

The mean potential scale reduction factor was 1.011 (95% CI: 0.998–1.072), indicating MCMC convergence. The overall model explained an average variation of 12.5% (Tjur *R^2^* = 0.125) across the 70 species (Fig. 3; Table S9). The fixed effects component (82.5%) explained 4.7 times more variance than the random effects component (spatial effects; 17.5%), highlighting that the predictors captured variation among sites (Fig. 3). Among the predictors, the proportion of native trees, a site-level predictor, accounted for the highest explained variation in species occurrences (35.1%), followed by cashew cover (20.2%), forest cover (15.1%), and canopy cover (12.1%) (Fig. 3). Thus, site-level predictors explained 47.2% and landscape-level predictors explained 35.3% of the variation in the data.

**Figure 3.**
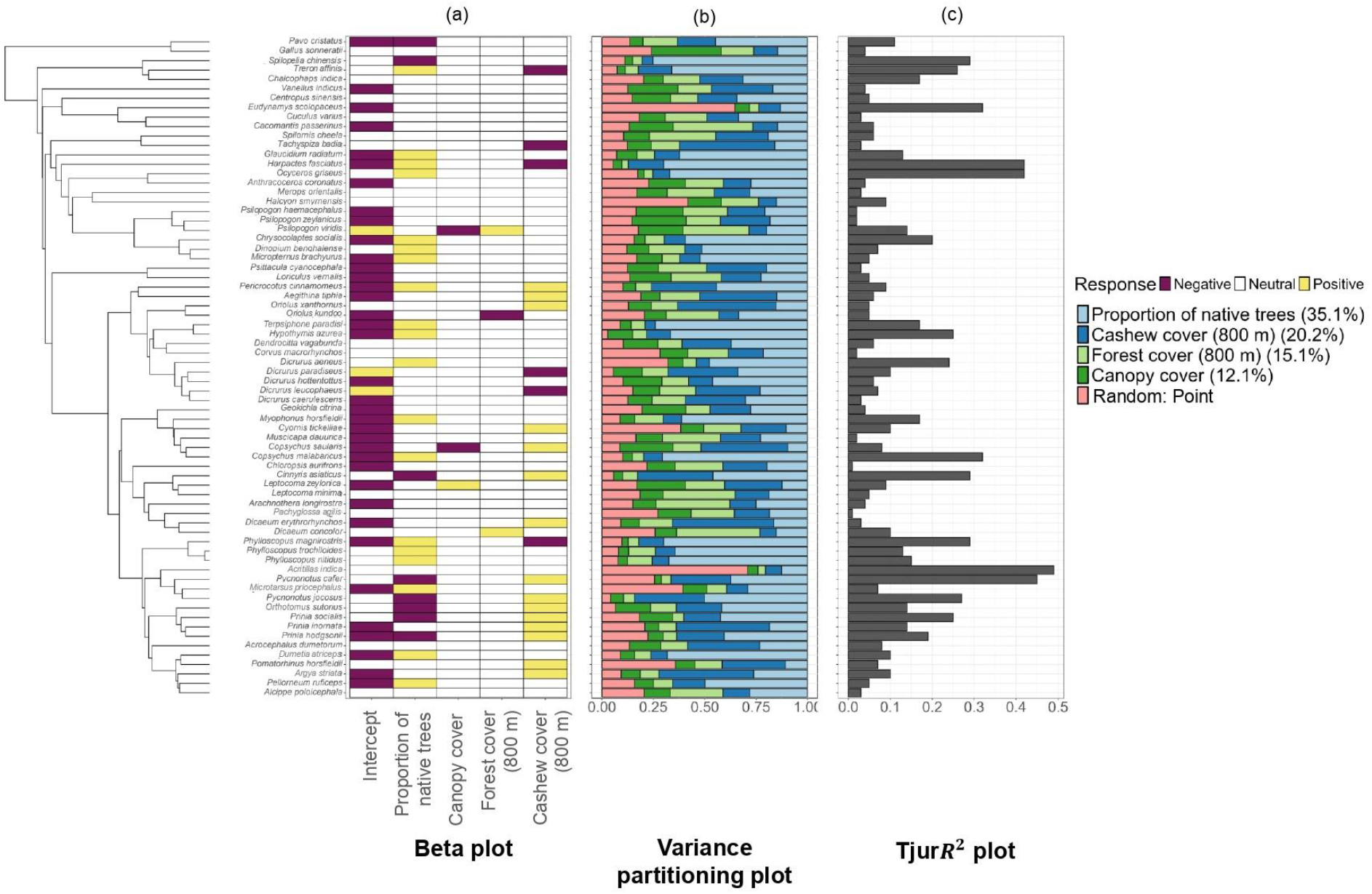
Composite figure showcasing the phylogenetic tree with tips representing bird species, (a) Beta plot (posterior regression parameters), (b) Variance partitioning plot and (c) Tjur*R^2^* plot. Beta plot displays responses of species to local- and landscape-level predictors. Yellow and violet boxes indicate statistically significantly positive and negative responses. Variance partitioning plot shows the proportion of variance in species occurrence attributed to different predictors. Tjur*R^2^* plot illustrates the variation in species occurrences explained by the overall model.

**Figure 4.**
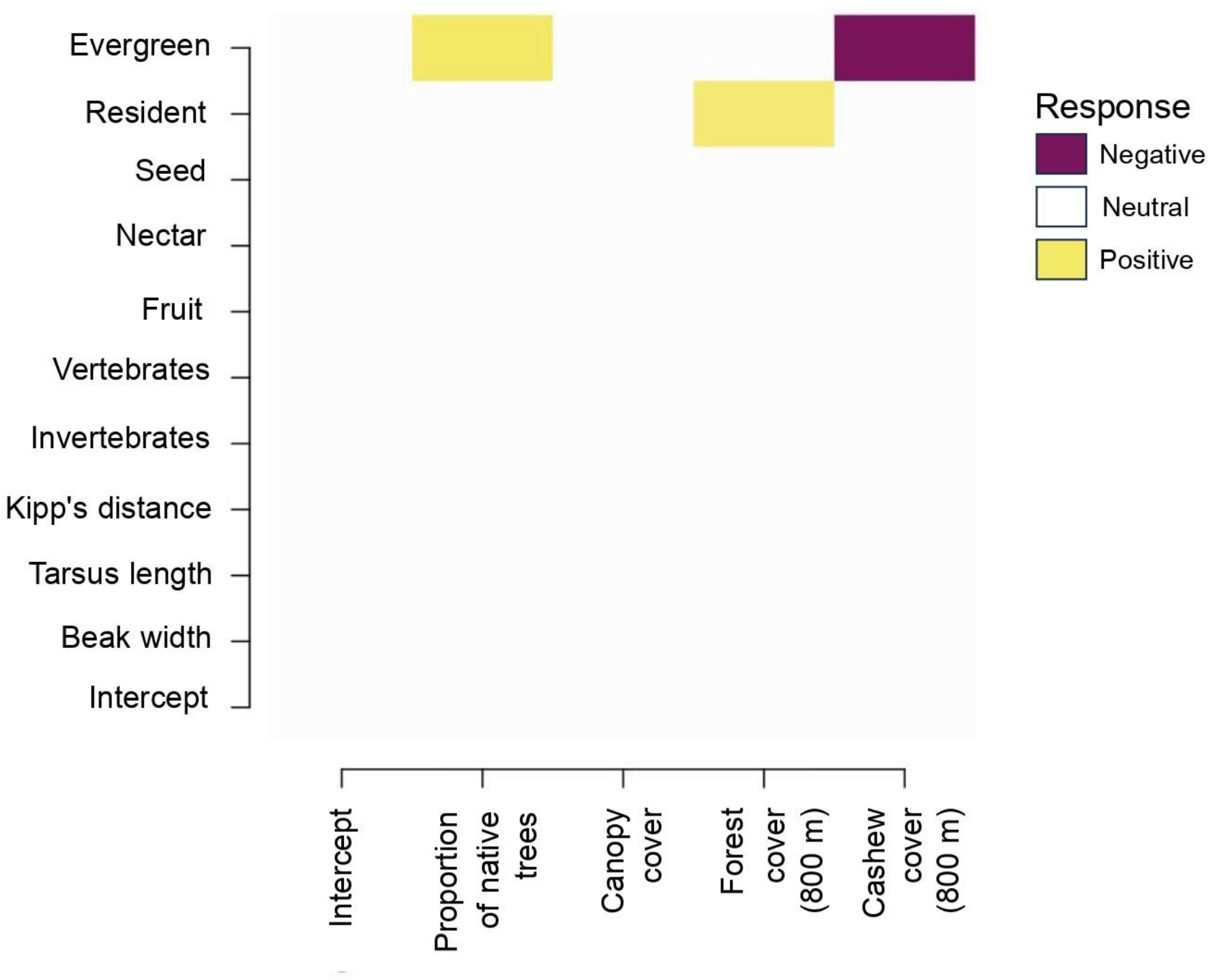
Gamma plot illustrating trait responses to local- (proportion of native trees and canopy cover) and landscape-level (forest cover and cashew cover) predictors. Yellow and violet boxes indicate statistically significantly positive and negative responses, respectively.

The beta coefficients for at least 30 species showed statistically significant effects for site-level variables (proportion of native trees: 27 species; canopy cover: three species), while 24 species showed statistically significant effects for landscape-level variables (forest cover: three species; cashew cover: 21 species) (Fig. 3). For 17 out of the 70 species, 50% or more of the variation in their occurrences was explained by the proportion of native trees alone (Table S9). The probability of occurrence increased for 19 species (e.g., *Ocyceros griseus*) (Fig. S6) and decreased for eight species (e.g., *Pavo cristatus*) (Fig. S5) with the proportion of native trees. The probability of occurrence decreased for two species (*Copsychus saularis* and *Psilopogon viridis*) (Fig. S7) and increased for one species (*Cinnyris asiaticus*) (Fig. S8) with canopy cover. The probability of occurrence decreased for one species (*Oriolus kundoo*) (Fig. S9) and increased for two species (*Dicaeum concolor* and *Psilopogon viridis*) (Fig. S10) with forest cover (Fig. S10). The probability of occurrence decreased for six species (e.g., *Harpactes fasciatus*) (Fig. S11) and increased for fifteen species (e.g., *Prinia* spp.) (Fig. S12) with cashew cover.

Traits explained 11% of the variation in species occurrences. The probability of occurrence for evergreen forest species was positively associated with the proportion of native trees and negatively with cashew cover (Fig. 5, Fig. S13). The probability of occurrence for resident species increased with increasing forest cover (Fig. 5 Fig. S13). We detected a phylogenetic signal in the residuals (⍴ = 0.59, 95% CI: 0.18–0.82), indicating that closely-related species were similar in their responses to predictor variables.

## 4. DISCUSSION

We tested the effects of site- and landscape-level factors on bird diversity, composition, traits and species occurrence by sampling across cashew intensification gradients from cashew agroforests with native trees to cashew monoculture plantations. At the community level, only the proportion of native trees positively influenced taxonomic and phylogenetic diversities. At the species level, along with this, cashew cover explained significant variation in species occurrences. Evergreen forest specialist species were influenced positively by the site-level predictor (proportion of native trees) and negatively by the landscape-level predictor (cashew cover). Resident species benefited from forest cover. This study demonstrates the importance of maintaining native trees in plantations and retaining forest patches in agricultural landscapes.

### Role of site and landscape-level factors

We demonstrate the critical role of native trees in conserving bird communities, deviating from Pérez et al., (2024), but aligning with Warren-Thomas et al., (2020). Cashew agroforests performed better than monocultures, underscoring the importance of retaining native trees within orchards, especially for evolutionarily distinct species with an affinity for evergreen forests. Since the proportion of native trees was strongly correlated with native tree species richness and forest structure variables, we can conclude that forest structural integrity and tree diversity are critical in influencing bird occurrence and diversity. In this landscape, where most land is privately owned and likely to be converted to monocultures, government policy is needed to ensure native trees are retained or planted within them. The effects of native tree cover on cashew yields need to be evaluated. If favourable, it can aid conservation.

Contrary to findings of Barbaro et al., (2021), none of the community-level metrics were associated with landscape-level variables. However, several species, especially specialists and residents, were associated with them. Cashew cover is increasing in the landscape (Rege and Lee, 2022). Along with retaining native trees, retaining forest cover in the landscape is critical. The government should protect existing Reserved Forest patches and restore degraded forest patches with appropriate native evergreen tree species to host evergreen forest bird species. This study recommends a hybrid approach combining sparing land for biodiversity and making agriculture more bird-friendly.

### Sharing-sparing implications

Our landscape features elements of both land-sharing and sparing— high-intensity cashew monocultures, low-intensity cashew agroforests, and forest patches (Sidemo Holm et al., 2021), similar to the ‘three-compartment strategy’ (Finch et al., 2019), that maximised bird populations in agricultural landscapes of England. Unfortunately, agroforests are declining as they are increasingly being replaced by monocultures (VS pers. obs.). Government-owned Reserved Forests are likely to remain relatively intact, provided they are not diverted for non- forestry uses. These changes are shifting the landscape from the current three-compartment strategy towards a land-sparing strategy. This may be detrimental to bird diversity, given the strong influence of site-level variables on bird diversity and prevalence.

In the case of neighbouring district Ratnagiri, with more than 70% of its geographic area under forests, approximately 97% of these is privately owned. The landscape could be converted into vast expanses of monocultures, like in states like Kerala in the southern Western Ghats (Balan et al., 2020). Conversion to plantations and forest fragmentation are ascribed as drivers of destructive landslides in Kerala (Vijayan et al., 2024), an aspect land- planners should consider for this landscape.

### Differential responses of diversity measures

While taxonomic and phylogenetic diversity increased with increasing proportion of native trees at a site, we did not find any pattern for functional diversity. Such non-congruent patterns among different diversity measures (Sreekar et al., 2021), highlight their complementary insights on bird persistence.

More than 27% of species showed positive associations with the proportion of native trees, compared to the 13% showing negative associations, demonstrating its positive influence on taxonomic diversity. Species benefiting from the proportion of native trees included lineages with long branch lengths, such as trogons, woodpeckers, and hornbills, explaining the positive association of phylogenetic diversity with it.

The absence of association between functional diversity and diet-related traits with predictors can be explained by ‘compensatory dynamics’, where specialists are replaced by generalists (Cooke et al., 2019). Forest specialist frugivores (e.g., *Ocyceros griseus*) and insectivores (Picidae members) were negatively impacted and replaced by their generalist counterparts (*Pycnonotus* spp., *Prinia* spp.).

Unlike in other studies (Biswas et al., 2023; Bonfim et al., 2024), we found a phylogenetic signal in species responses. *Phylloscopus* warblers and *Prinia* spp. exhibited positive and negative associations with native tree cover.

### Different responses at community- and species-level

Multiple conservation goals exist, especially in wet tropical mountain ranges, the most threatened habitats harbouring small-ranged species (Orme et al., 2006). Maximising bird diversity should not come at the expense of species of conservation concern. An examination solely at the community level (Chapman et al., 2018), would not have highlighted the negative influence of cashew cover on forest specialists, highlighting the need to examine bird responses from species and community levels simultaneously to inform conservation better.

### 4.3. Conservation Implications

We demonstrate the importance of both preserving existing forest patches amidst monocultures and retaining native trees within them to conserve avian diversity and species of conservation concern, consistent with recommendations made for rubber and coffee plantations globally (Alvarez-Alvarez et al., 2022; Warren-Thomas et al., 2020). Given the rapid expansion of cashew monocultures in other tropical regions globally (Powell et al., 2023; Rege et al., 2024), policies should encourage large landowners to retain forest patches and plant native trees with high functional value to enhance bird diversity.

Privately-owned forest patches not converted to plantations are repeatedly cleared (after an interval of 8-10 years) for fuelwood, shifting the forest type from evergreen to deciduous (Biswas et al., 2024). Landowners should be incentivised to restore degraded deciduous forests to benefit forest specialists of conservation concern.

We advocate simultaneously examining community, species, and trait level responses to human activities as they provide complementary insights. Studies should investigate the influence of native tree cover and proximity to forests on pollination and herbivory in plantations to inform land-management policy in human-dominated landscapes.

## ACKNOWLEDGEMENTS

We thank the Dean and Director of the Wildlife Institute of India and the Maharashtra Forest Department (No: Desk-22(8)/WL/Research/C.No. 103/635/24-25) for their support. We also acknowledge the financial assistance provided by the Wildlife Institute of India, Godrej Consumer Products Limited and Rohini Nilekani Philanthropies. Special thanks to Pravin, Shital Desai, and Himanshu Lad for their invaluable support during fieldwork. Thanks are also due to Arpitha Jayanth, Jithin Vijayan, and Sanath Manimoole for their help with analyses, and to Jahnavi Joshi for extracting the phylogenetic trees. NM and RN thank Navendu Page for his invaluable inputs and help with plant identification. RN thanks Anand Osuri and Kulbhushansingh Suryawanshi for insightful discussions.

## AUTHORS’ CONTRIBUTIONS

Conceptualisation: RN, NM, RJ; Data curation: NM, VS; Formal analysis: NM, RN, NB; Funding acquisition: NM, RN; Investigation: NM, VS; Methodology: RN, NM, RJ; Project administration: VS, NM; Resources: NM, RN; Supervision: RN, RJ; Visualisation: RN; Writing – original draft: NM, RN; Writing – review & editing: AR, RJ, NB,VS. All authors in this study are indigenous to the country where the study was conducted. Whenever relevant, literature published by scientists from the region has been cited. All authors gave final approval for publication.

## DATA ACCESSIBILITY STATEMENT

The associated data and R-codes will be uploaded on Zenodo (https://zenodo.org) upon acceptance of this manuscript.

## SUPPLEMENTARY INFORMATION

### Supplementary Images

**Figure S1.**
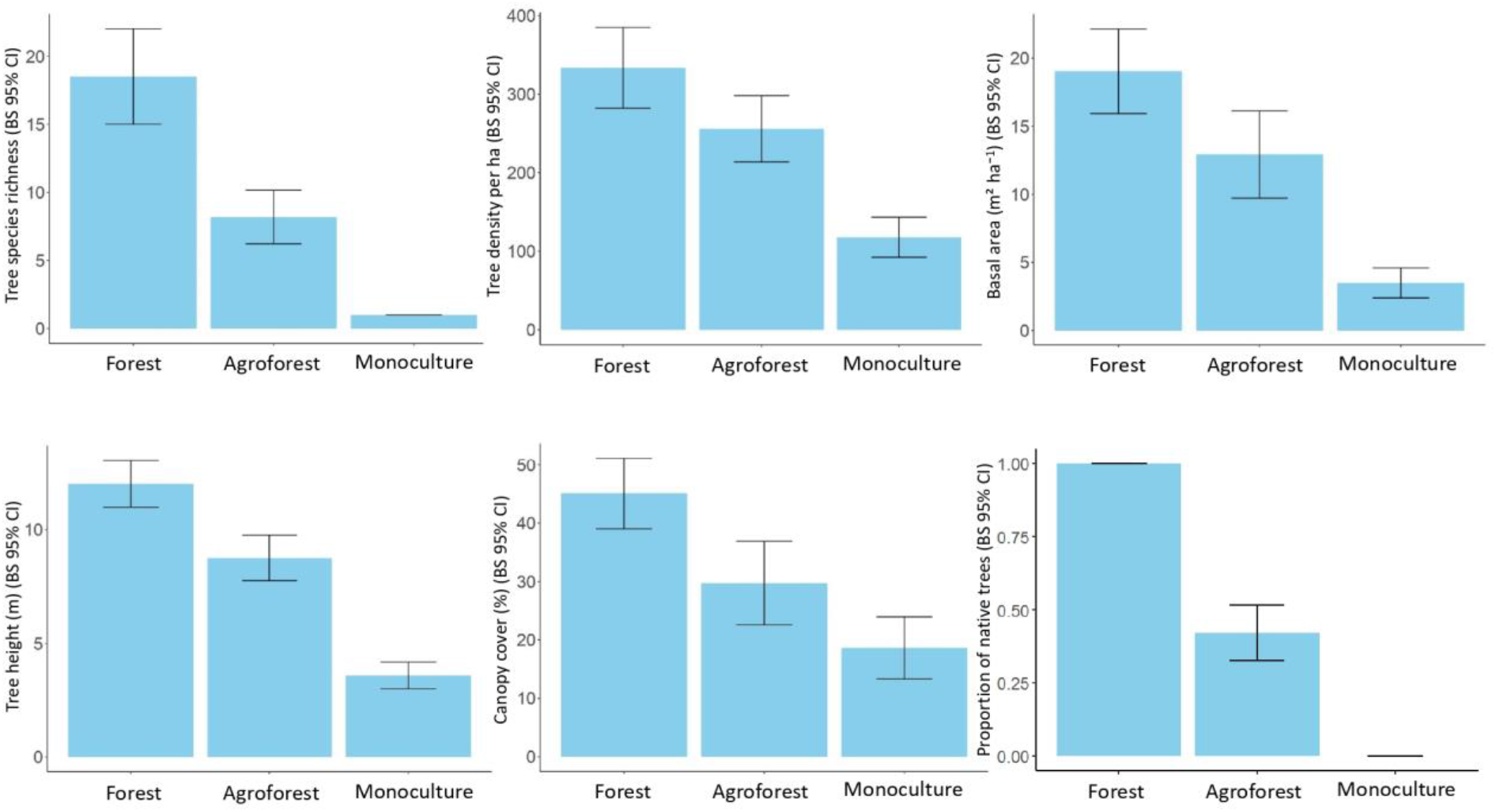
Bar plots comparing tree species richness, vegetation structural attributes, and proportions of native and cashew trees across the land-use categories. Error bars represent bootstrapped (n = 1000) 95% CIs.

**Figure S2.**
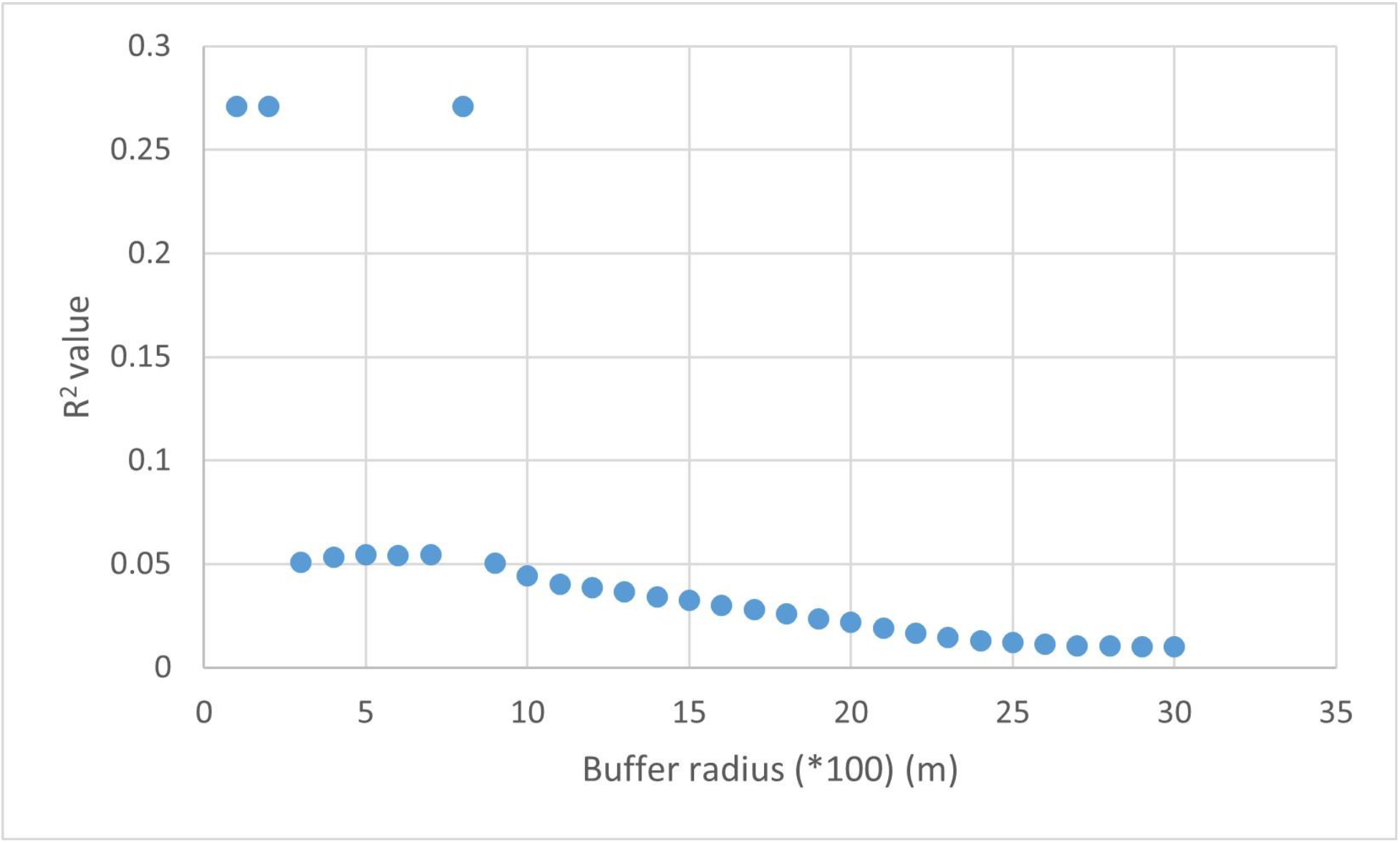
*R^2^* values for models testing for association between species richness and amount of forest cover at different buffer radii (100 m to 3000 m around each sampling point). We did this to determine the optimum buffer radius for landscape-level variables following the scale-of-effect approach suggested by Fahrig, (2013).

**Figure S3.**
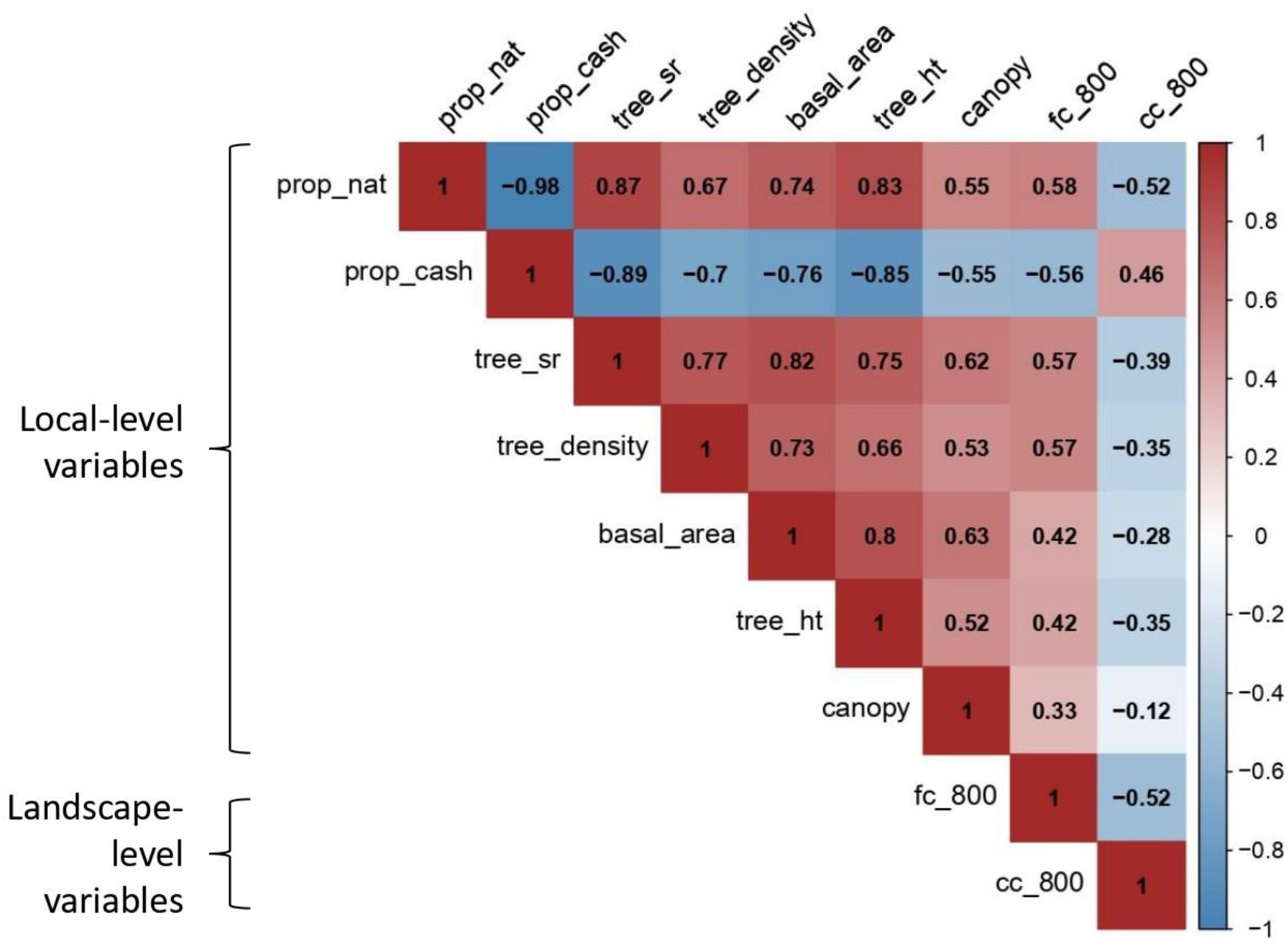
Correlation biplot depicting correlations among local and landscape predictors. Values inside boxes represent Pearson’s correlation coefficients. The scale indicates a positive to negative gradient in correlation coefficients. prop_nat: proportion of native trees per plot, prop_cash: proportion of cashew trees per plot, tree_sr = tree species richness per plot, tree_density: tree density per hectare, basal area: basal area (m^2^ ha^-1^), tree_ht: tree height (m), canopy: canopy cover (%), fc_800: proportion of forest cover in 800 m buffer radius, cc_800: proportion of cashew cover in 800 m buffer radius, Proportions of forest and cashew cover around each point were extracted from an existing land-use land-cover map (Rege et al. 2022).

**Figure S4.**
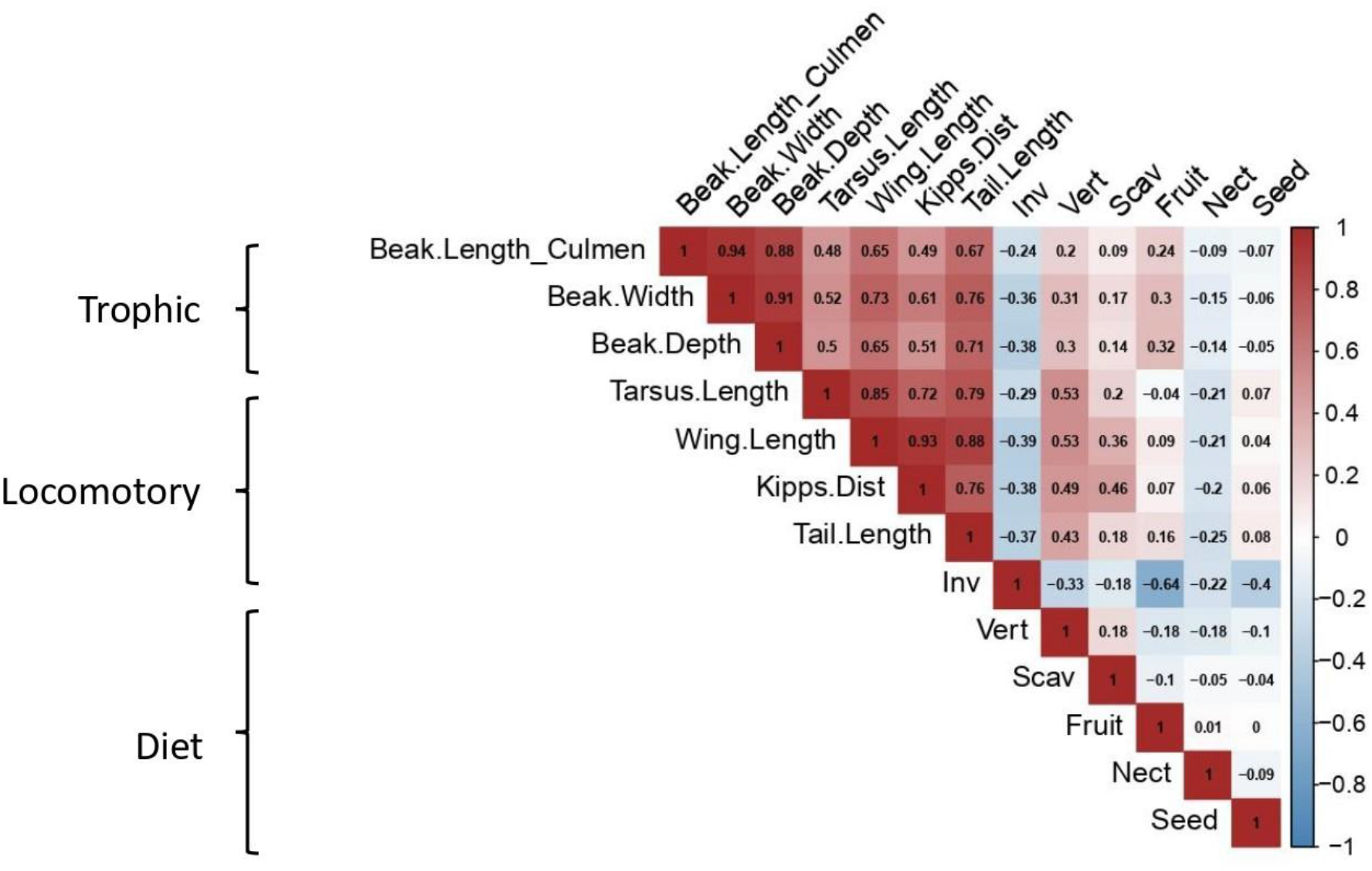
Correlation biplot depicting correlations among traits. Values inside boxes represent Pearson’s correlation coefficients. The scale indicates a positive to negative gradient in correlation coefficients. Inv: percentage invertebrates in diet, Vert: percentage vertebrates in diet, Scav: percentage scavenging, Fruit: percentage fruit in the diet, Nect: percentage nectar in the diet, Seed: percentage of seeds in the diet. The morphological traits were sourced from AVONET (Tobias et al., 2022), dietary traits were sourced from (Wilman et al., 2014).

**Figure S5.**
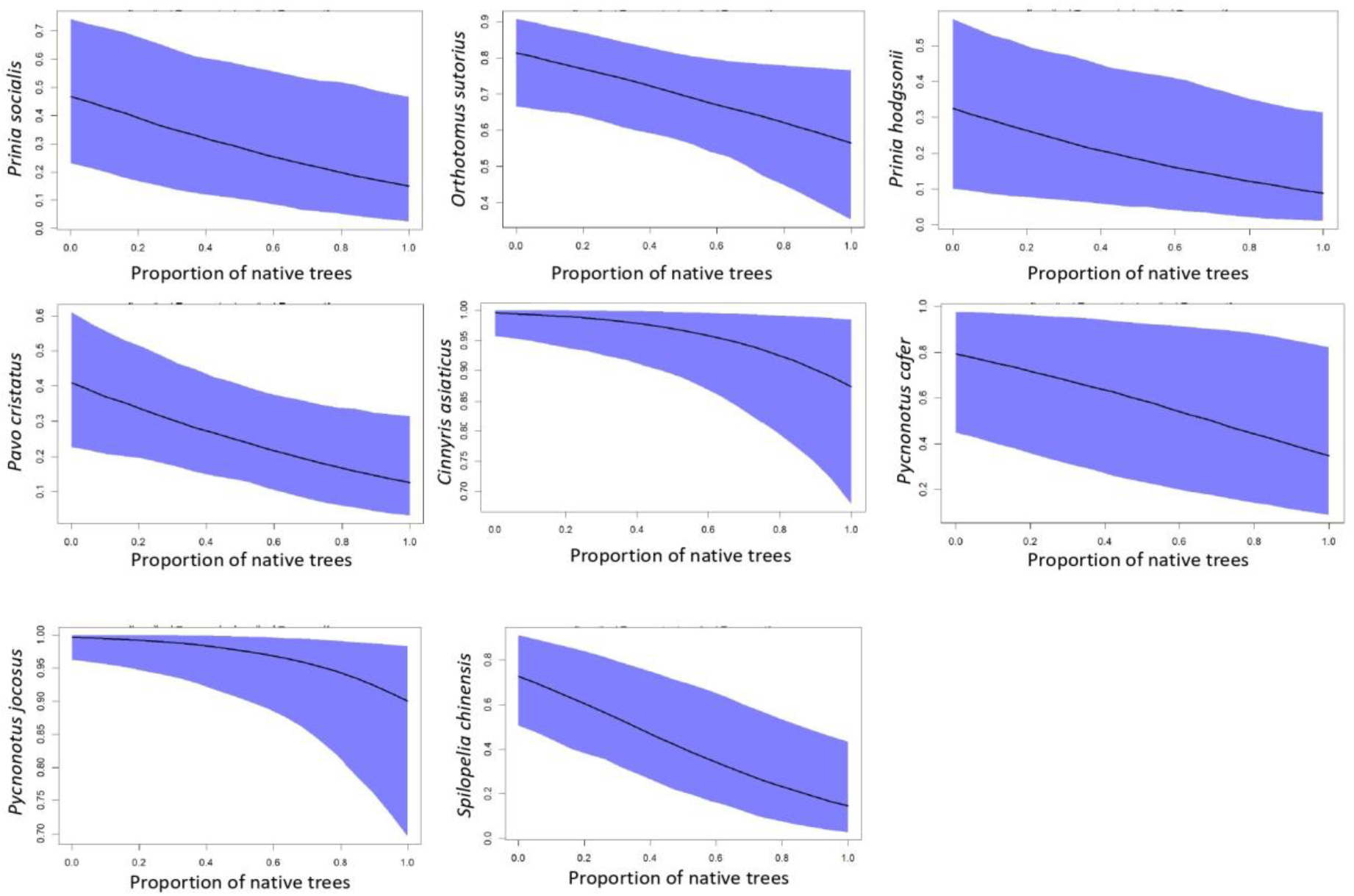
Predicted probability of occurrence of bird species (see y-axis labels for species ID) along a gradient of proportion of native trees. These are the species whose probabilities of occurrence decrease with increasing proportion of native trees.

**Figure S6.**
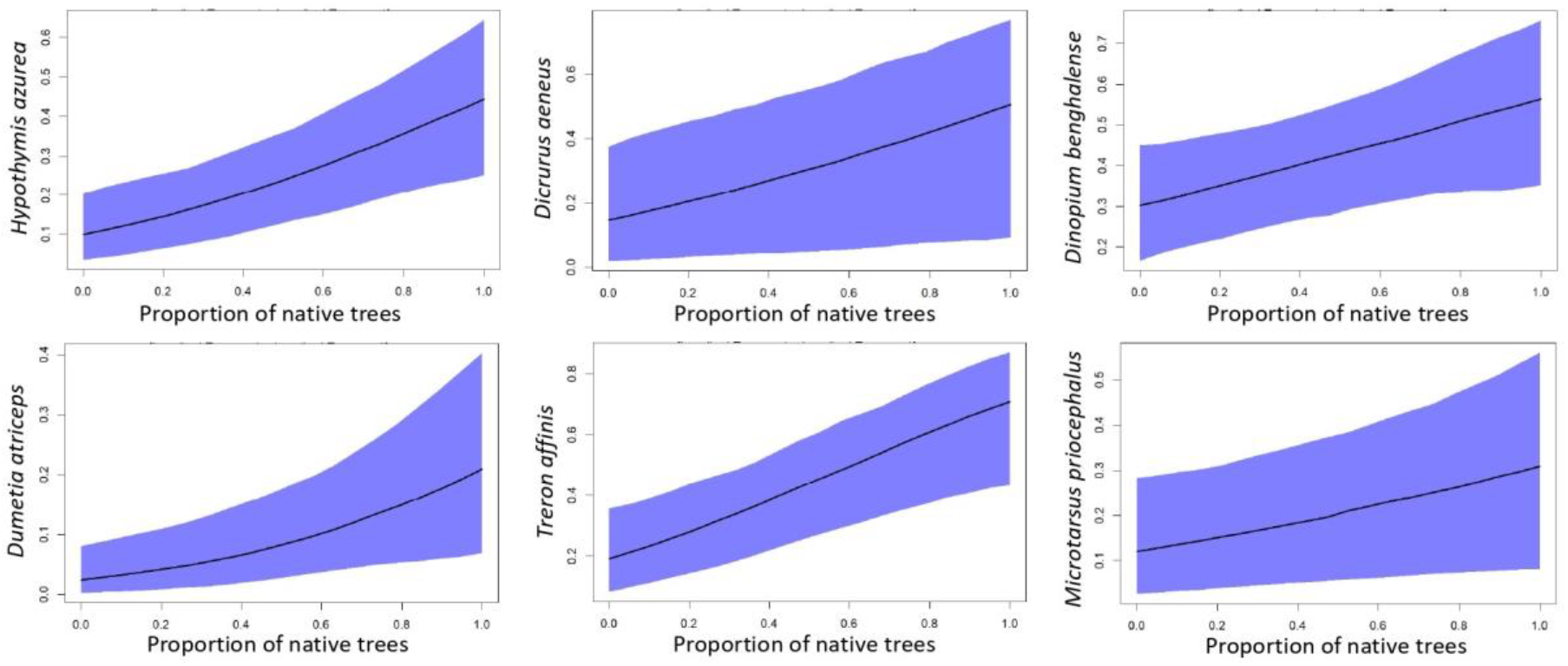
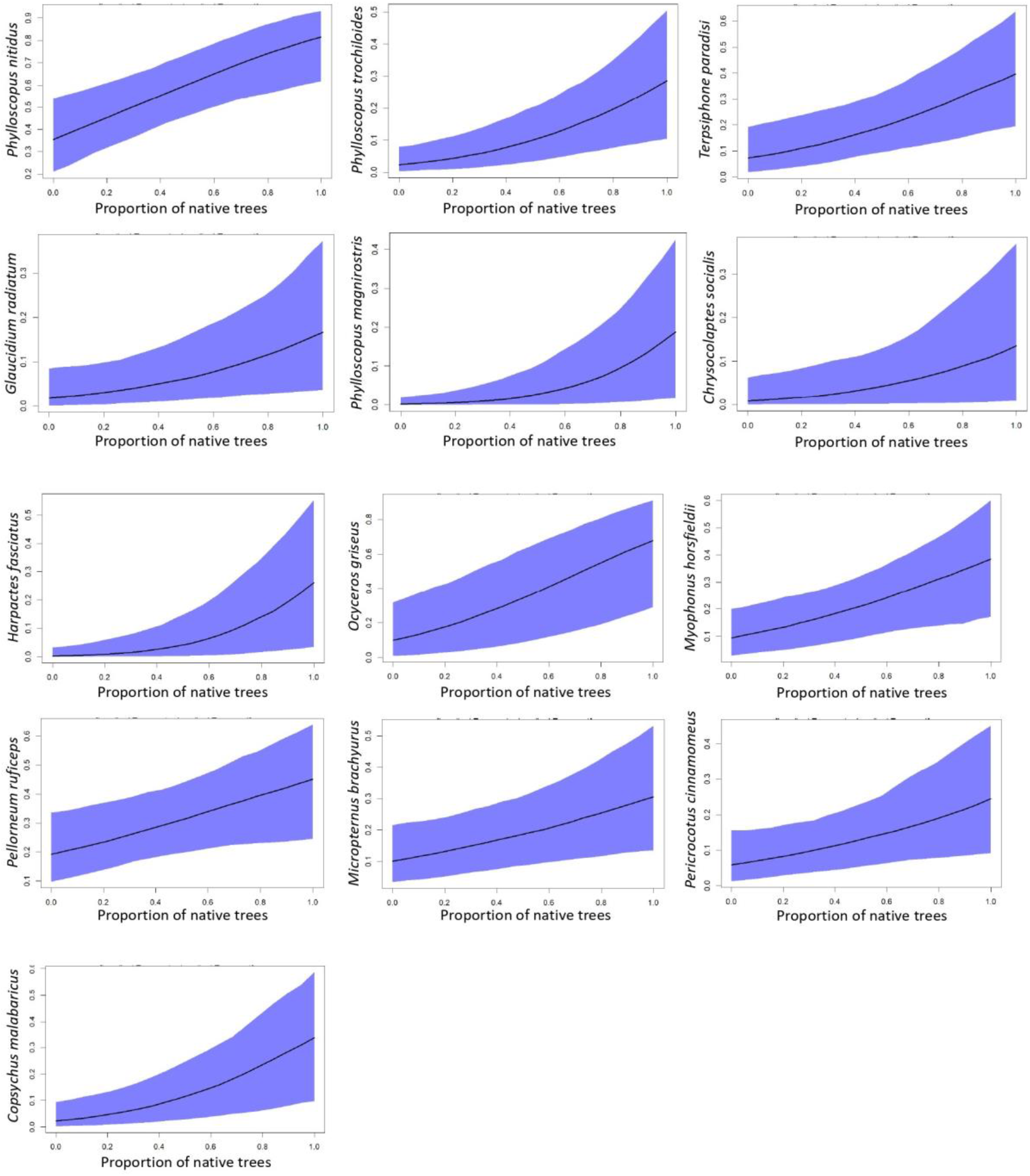
Predicted probability of occurrence of bird species (see y-axis labels for species ID) along a gradient of proportion of native trees. These are the species whose probabilities of occurrence increase with increasing proportion of native trees.

**Figure S7.**
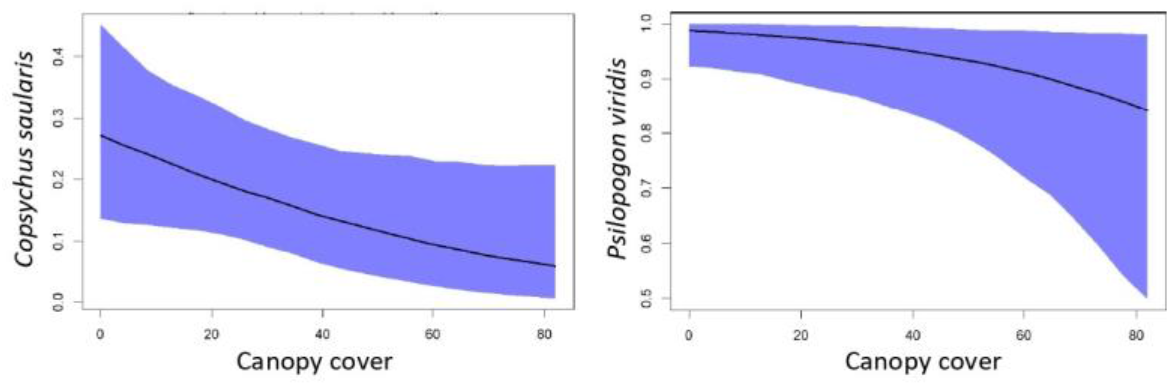
Predicted probability of occurrence of bird species (see y-axis labels for species ID) along a gradient of canopy cover. These are the species whose probabilities of occurrence decrease with increasing canopy cover.

**Figure S8.**
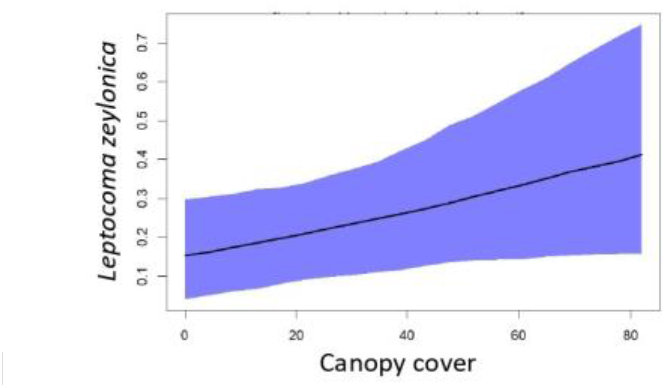
Predicted probability of occurrence of Purple-rumped Sunbird *Leptocoma zeylonica* along a gradient of canopy cover. This species has a higher probability of occurrence associated with increasing canopy cover.

**Figure S9.**
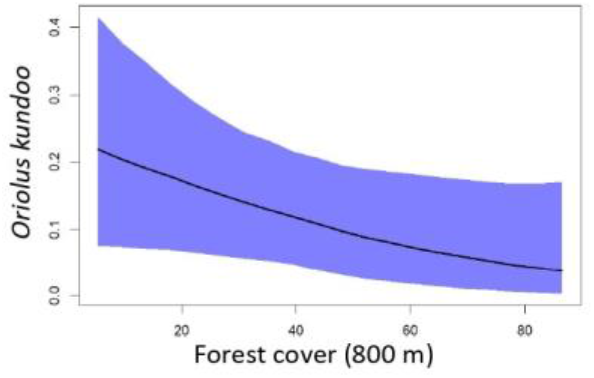
Predicted probability of occurrence of Indian Golden Oriole *Oriolus kundoo* along a gradient of forest cover in 800 m radius. This species has a lower probability of occurrence associated with increasing forest cover in 800 m radius.

**Figure S10.**
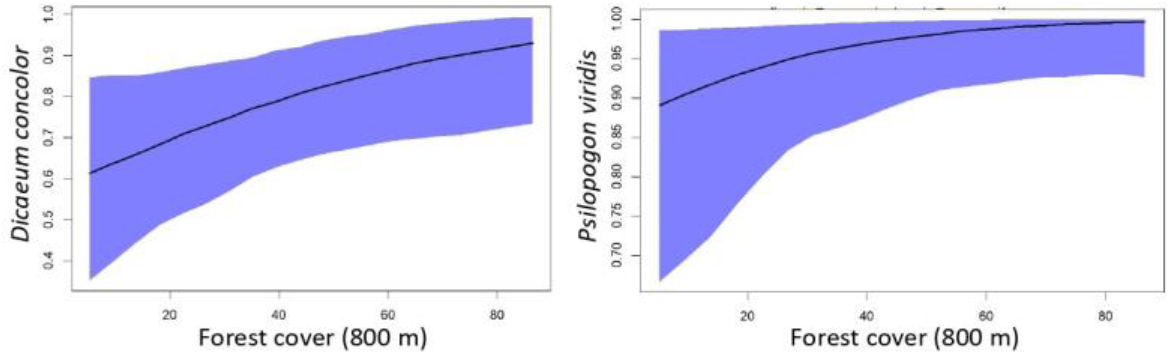
Predicted probability of occurrence of bird species (see y-axis labels for species ID) along a gradient of forest cover in 800 m radius. These are the species whose probabilities of occurrence increase with increasing forest cover in 800 m radius.

**Figure S11.**
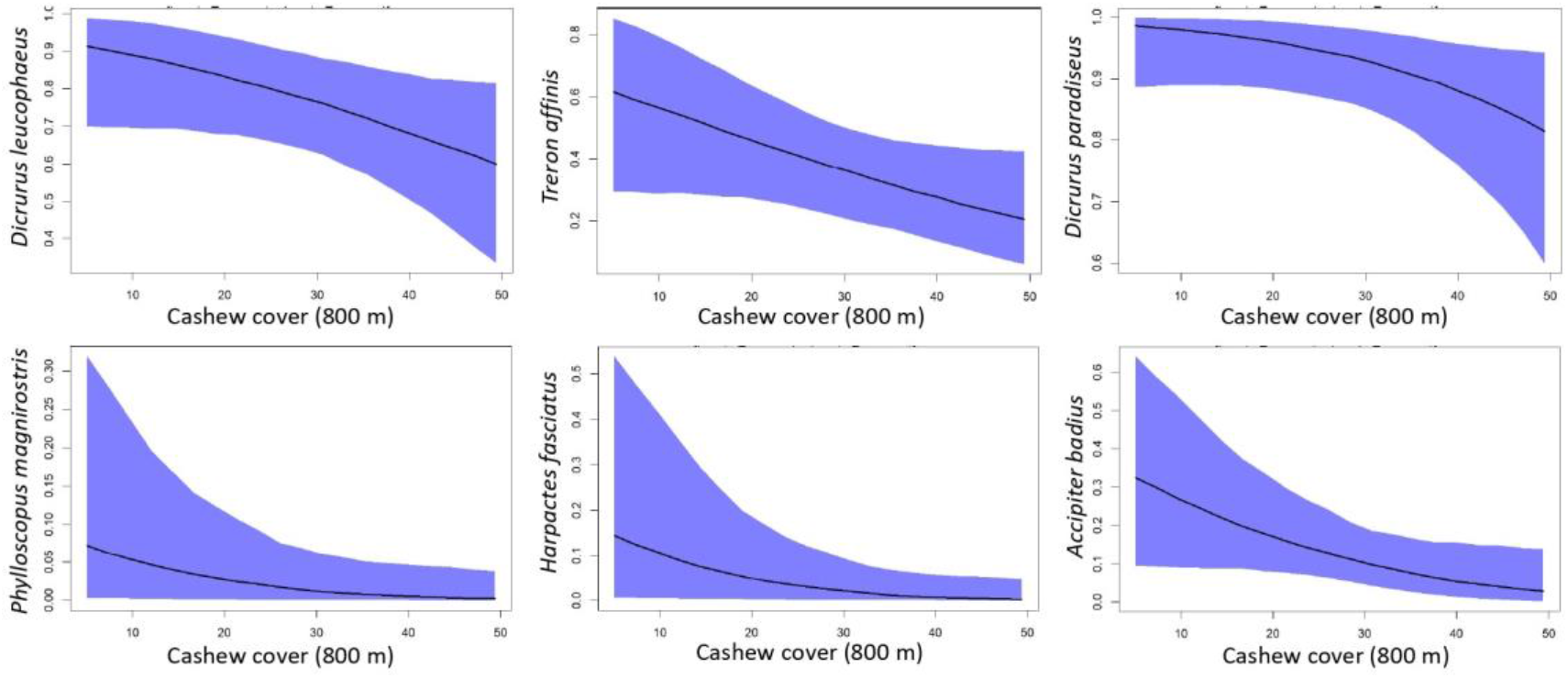
Predicted probability of occurrence of bird species (see y-axis labels for species ID) along a gradient of cashew cover in 800 m radius. These are the species whose probabilities of occurrence decrease with increasing cashew cover in 800 m radius.

**Figure S12.**
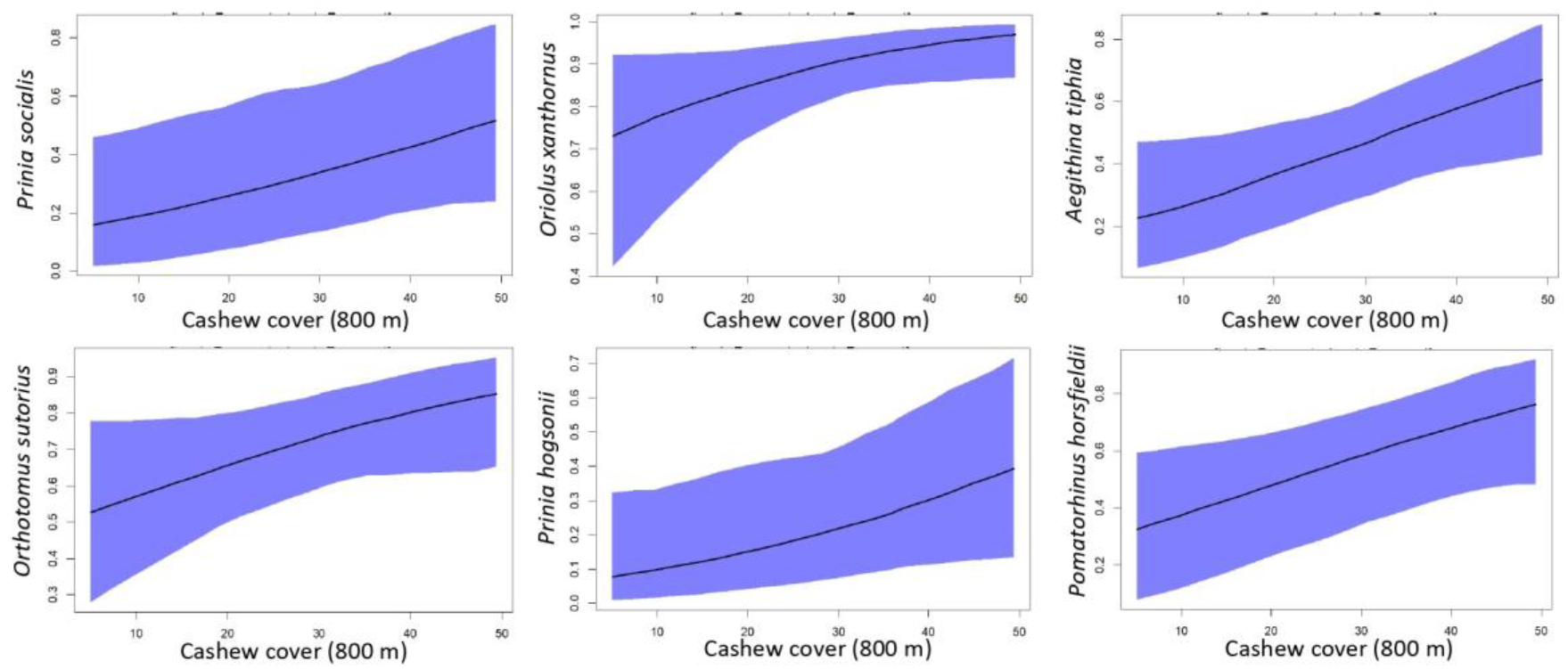
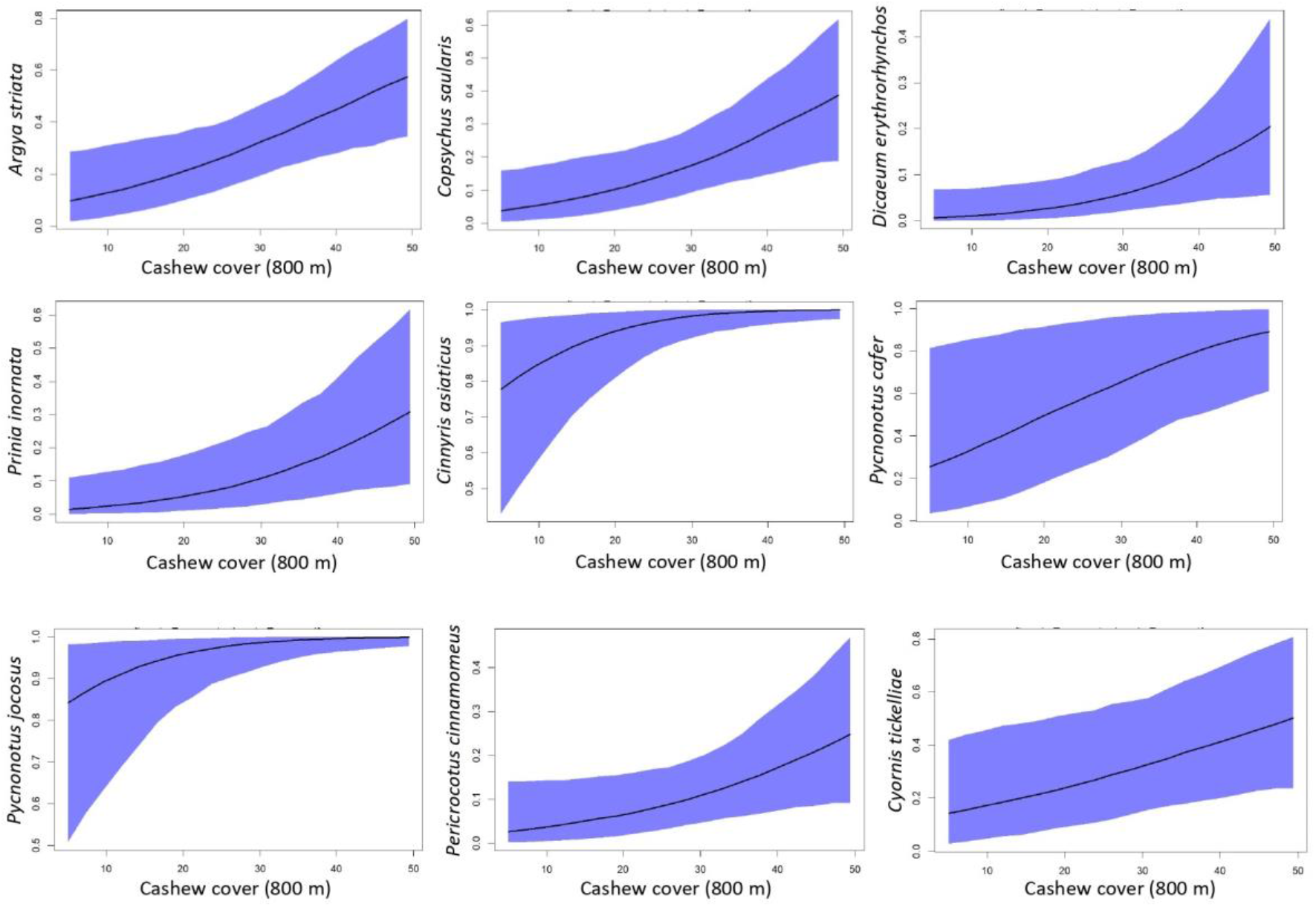
Predicted probability of occurrence of bird species (see y-axis labels for species ID) along a gradient of cashew cover in 800 m radius. These are the species whose probabilities of occurrence increase with increasing cashew cover in 800 m radius.

**Figure S13.**
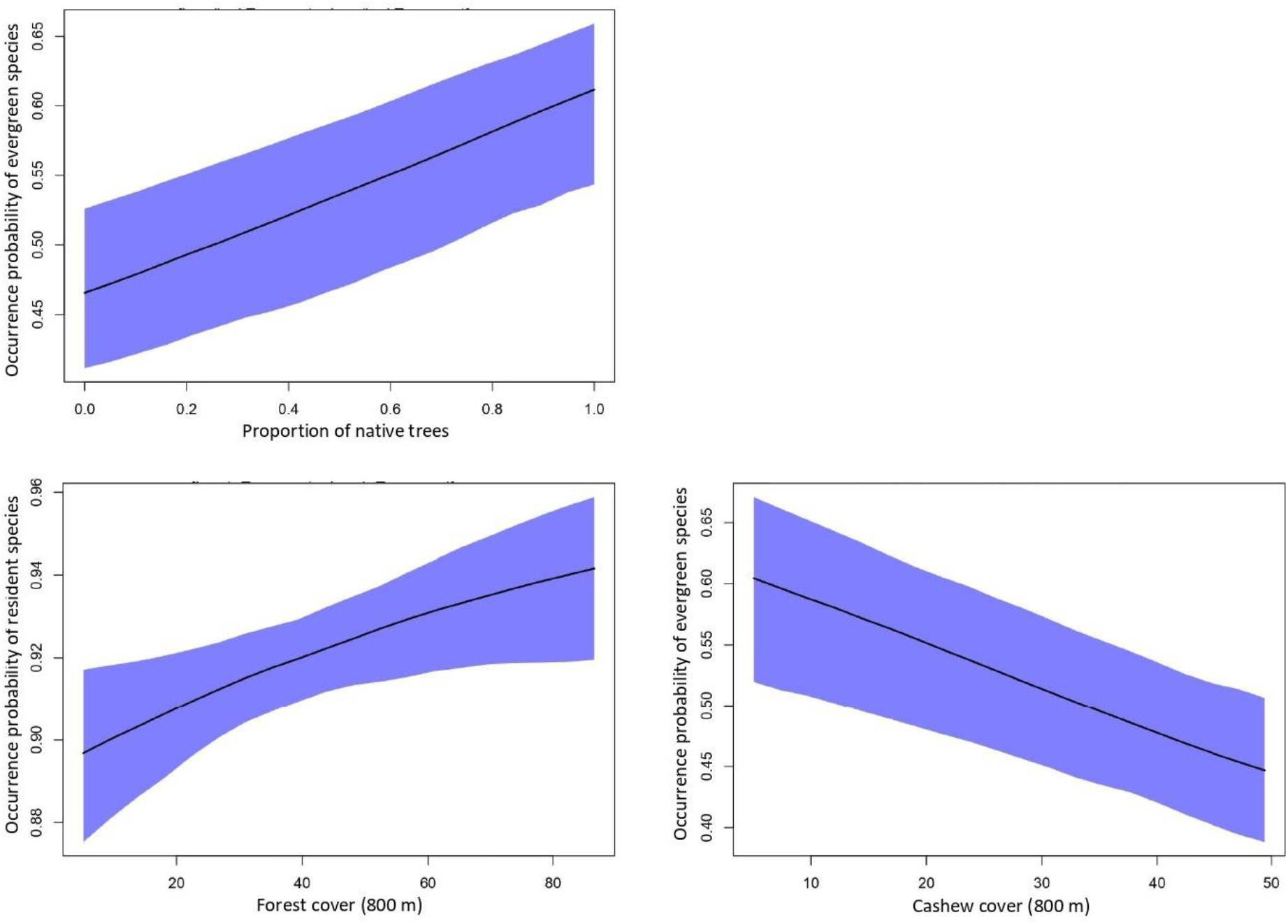
Probabilities of occurrence of evergreen and resident species along gradients of proportion of native trees, forest cover in 800 m buffer, and cashew cover in 800 m buffer.

**Table S1.** Checklist of tree species detected in 100 plots of a 10 m circular radius. We also report the number of plots where each species was detected and the total number of individuals detected. Species are sorted according to families.

| No. | Family | Species | Total No.<br>across<br>habitats | No.of plots in<br>which the<br>species was<br>recorded |
| --- | --- | --- | --- | --- |
| 1 | Anacardiaceae | *Anacardium occidentale* | 272 | 66 |
| 2 | Anacardiaceae | *Holigarna arnottiana* | 16 | 5 |
| 3 | Anacardiaceae | *Holigarna grahamii* | 1 | 1 |
| 4 | Anacardiaceae | *Mangifera indica* | 2 | 2 |
| 5 | Anacardiaceae | *Nothopegia castaneifolia* | 1 | 1 |
| 6 | Annonaceae | *Miliusa tomentosa* | 4 | 4 |
| 7 | Annonaceae | *Monoon fragrans* | 1 | 1 |
| 8 | Apocynaceae | *Alstonia scholaris* | 2 | 2 |
| 9 | Apocynaceae | *Tabernaemontana*<br>*alternifolia* | 8 | 6 |
| 10 | Arecaceae | *Caryota urens* | 1 | 1 |
| 11 | Arecaceae | *Cocos nucifera* | 2 | 1 |
| 12 | Bignoniaceae | *Stereospermum*<br>*angustifolium* | 1 | 1 |
| 13 | Celastraceae | *Lophopetalum*<br>*wightianum* | 1 | 1 |
| 14 | Clusiaceae | *Garcinia indica* | 3 | 3 |
| 15 | Combretaceae | *Terminalia bellirica* | 4 | 4 |
| 16 | Combretaceae | *Terminalia elliptica* | 21 | 12 |
| 17 | Combretaceae | *Terminalia paniculata* | 80 | 33 |
| 18 | Dilleniaceae | *Dillenia pentagyna* | 5 | 5 |
| 19 | Ebenaceae | *Diospyros candolleana* | 7 | 3 |
| 20 | Elaeocarpaceae | *Elaeocarpus*<br>*tuberculatus* | 1 | 1 |
| 21 | Euphorbiaceae | *Macaranga peltata* | 1 | 1 |
| 22 | Euphorbiaceae | *Mallotus philippensis* | 3 | 2 |
| 23 | Fabaceae | *Acacia auriculiformis* | 11 | 4 |
| 24 | Fabaceae | *Dalbergia sissoo* | 1 | 1 |
| 25 | Fabaceae | *Saraca asoca* | 1 | 1 |
| 26 | Fabaceae | *Xylia xylocarpa* | 22 | 12 |
| 27 | Lamiaceae | *Tectona grandis* | 10 | 4 |
| 28 | Lamiaceae | *Vitex altissima* | 2 | 2 |
| 29 | Lauraceae | *Machilus glaucescens* | 1 | 1 |
| 30 | Lecythidaceae | *Careya arborea* | 14 | 9 |
| 31 | Loganiaceae | *Strychnos nux-vomica* | 2 | 2 |
| 32 | Lythraceae | *Lagerstroemia*<br>*microcarpa* | 4 | 3 |
| 33 | Lythraceae | *Lagerstroemia parviflora* | 2 | 2 |
| 34 | Malvaceae | *Bombax ceiba* | 1 | 1 |
| 35 | Malvaceae | *Grewia tiliifolia* | 2 | 2 |
| 36 | Malvaceae | *Pterospermum*<br>*angustifolium* | 1 | 1 |
| 37 | Malvaceae | *Pterospermum*<br>*diversifolium* | 1 | 1 |
| 38 | Malvaceae | *Sterculia guttata* | 1 | 1 |
| 39 | Moraceae | *Artocarpus heterophyllus* | 4 | 3 |
| 40 | Moraceae | *Artocarpus hirsutus* | 1 | 1 |
| 41 | Myrtaceae | *Syzygium cumini* | 16 | 10 |
| 42 | Oleaceae | *Chionanthus mala-elengi* | 1 | 1 |
| 43 | Oleaceae | *Tetrapilus dioicus* | 5 | 4 |
| 44 | Phyllanthaceae | *Aporosa cardiosperma* | 39 | 13 |
| 45 | Phyllanthaceae | *Bridelia retusa* | 4 | 4 |
| 46 | Phyllanthaceae | *Phyllanthus emblica* | 1 | 1 |
| 47 | Rhizophoraceae | *Carallia brachiata* | 5 | 4 |
| 48 | Rubiaceae | *Adina cordifolia* | 1 | 1 |
| 49 | Rubiaceae | *Ixora brachiata* | 16 | 8 |
| 50 | Rubiaceae | *Meyna laxiflora* | 1 | 1 |
| 51 | Rubiaceae | *Mitragyna parvifolia* | 1 | 1 |
| 52 | Rutaceae | *Glycosmis*<br>*cochinchinensis* | 4 | 2 |
| 53 | Rutaceae | *Zanthoxylum rhetsa* | 2 | 2 |
| 54 | Salicaceae | *Flacourtia montana* | 5 | 4 |
| 55 | Sapindaceae | *Schleichera oleosa* | 9 | 6 |
| 56 | Sapotaceae | *Xantolis tomentosa* | 9 | 4 |

**Table S2.**
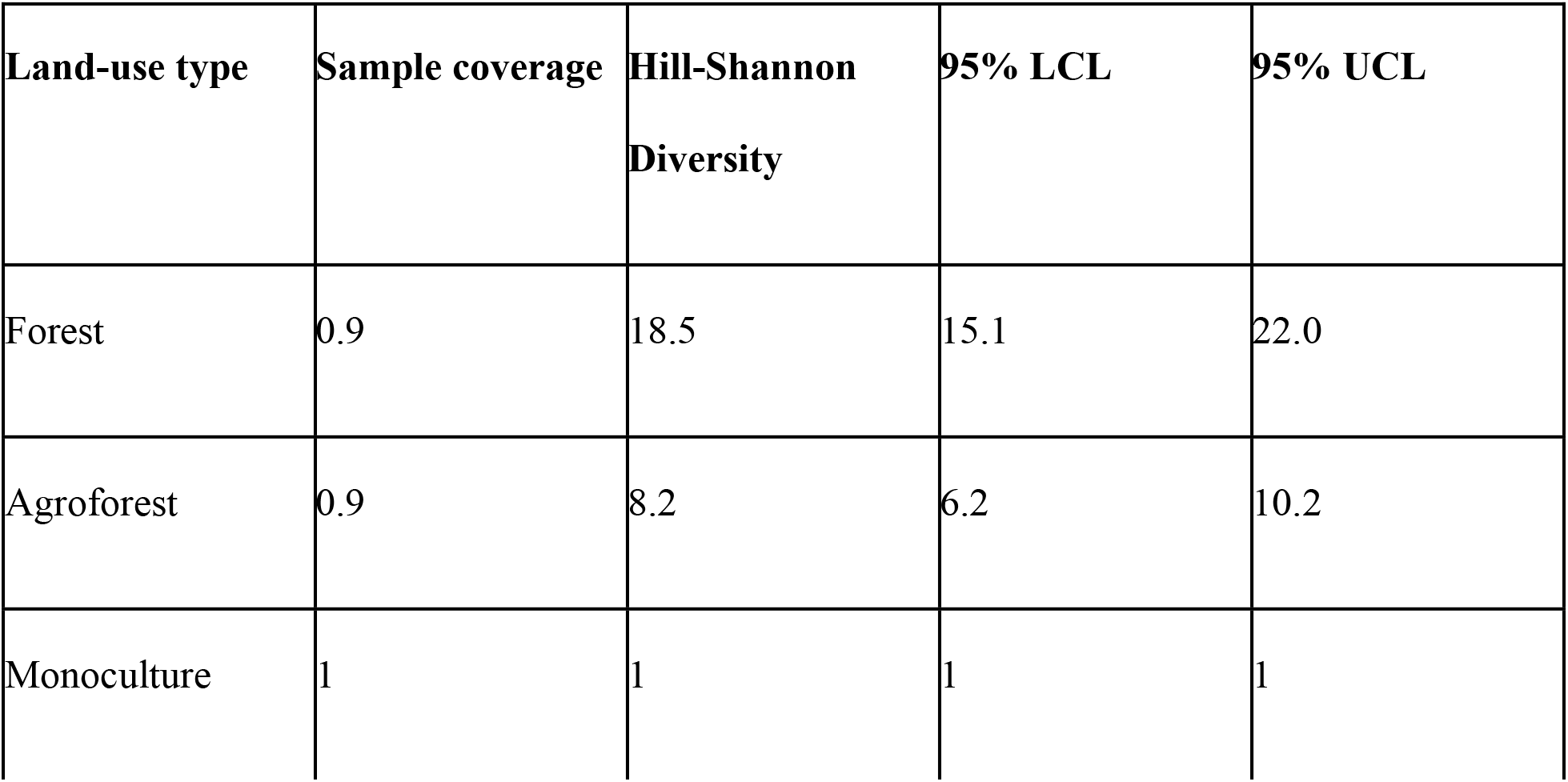
Summary of the sample coverage, Hill-Shannon diversity numbers, and the associated bootstrapped 95% CI of trees for the three land-use categories (forest, agroforest, and monoculture).

**Table S3.** Treatment contrast table comparing different site-level structural attributes across the three land-use categories (forests, cashew agroforest, and cashew monoculture).

| Structural attribute | Land-use type | Estimate | 95% CI | *p* |
| --- | --- | --- | --- | --- |
| Tree density per hectare<br>(natural log-transformed) | Intercept: Forest | 5.7305 | 5.527 – 5.933 | < **0.001** |
|  | Agroforest | -0.2915 | -0.575 – -0.007 | 0.0574 |
|  | Monoculture | -0.7713 | -1.041 – -0.501 | < **0.001** |
| Total basal area (m<sup>2</sup> ha<sup>-1</sup>) (natural log-transformed) | Intercept: Forest | 2.8395 | 2.475 – 3.203 | < **0.001** |
|  | Agroforest | -0.6025 | -1.111 – -0.093 | **0.0209** |
|  | Monoculture | -1.7363 | -2.220 – -1.252 | < **0.001** |
| Tree height (m) (natural log-transformed) | Intercept: Forest | 2.46789 | 2.350 – 2.585 | < **0.001** |
|  | Agroforest | -0.34782 | -0.512 – -0.183 | < **0.001** |
|  | Monoculture | -1.03301 | -1.189 – -0.876 | < **0.001** |
| Canopy cover (%) (natural log-transformed) | Intercept: Forest | 3.7594 | 3.481 – 4.038 | < **0.001** |
|  | Agroforest | -0.6104 | -1.001 – -0.220 | **0.003** |
|  | Monoculture | -0.6754 | -1.046 – -0.304 | < **0.001** |

**Table S4.** Tukey’s post-hoc results for vegetation structural attributes across land-use types.

| **Structural attribute** | **Contrast** | **Difference** | **95% LCI** | **95% UCI** | **p-adjusted** |
| --- | --- | --- | --- | --- | --- |
| Tree density per hectare | Agroforest - Forest | -0.292 | -0.632 | 0.049 | 0.109 |
|  | Monoculture -<br>Forest | -0.771 | -1.096 | -0.447 | **<0.001** |
|  | Monoculture -<br>Agroforest | -0.498 | -0.800 | -0.159 | **0.002** |
| Basal area (m<sup>2</sup><br>ha<sup>-1</sup>) | Agroforest -<br>Forest | -0.603 | -1.213 | 0.008 | 0.054 |
|  | Monoculture -<br>Forest | -1.736 | -2.318 | -1.155 | **<0.001** |
|  | Monoculture -<br>Agroforest | -1.134 | -1.708 | -0.559 | **<0.001** |
| Tree height (m) | Agroforest -<br>Forest | -0.348 | -0.546 | -0.150 | **<0.001** |
|  | Monoculture -<br>Forest | -1.033 | -1.221 | -0.845 | **<0.001** |
|  | Monoculture -<br>Agroforest | -0.685 | -0.871 | -0.499 | **<0.001** |
| Canopy cover<br>(%) | Agroforest -<br>Forest | -0.610 | -1.079 | -0.142 | **0.007** |
|  | Monoculture -<br>Forest | -0.675 | -1.121 | -0.230 | **0.002** |
|  | Monoculture -<br>Agroforest | -0.065 | -0.505 | 0.376 | 0.934 |

**Table S5.** Treatment contrast table comparing beta regression coefficients of proportion of native trees and proportion of cashew trees per plot across the three land-use categories (forest, agroforest, and monoculture).

| Attribute | Land-use type | Estimate | 95% CI | *p* |
| --- | --- | --- | --- | --- |
| Proportion of native trees | Intercept: Forest | 3.128 | 1.00 - 1.00 | **< 0.001** |
|  | Agroforest | -3.769 | 0.326 - 0.515 | **< 0.001** |
|  | Monoculture | -6.256 | 0.00 - 0.00 | **< 0.001** |

**Table S6.** Post-hoc results of Estimated Marginal Means for proportion of native trees and proportion of cashew trees per plot across land-use categories (forest, agroforest, and monoculture), using Tukey’s method of *p* value adjustment.

| Attribute | Contrast | Difference | 95%<br>LCI | 95%<br>UCI | $p$-adjusted |
| --- | --- | --- | --- | --- | --- |
| Proportion of native trees | Forest - Agroforest | 0.613 | 0.518 | 0.708 | <0.001 |
|  | Forest - Monoculture | 0.916 | 0.882 | 0.949 | <0.001 |
|  | Agroforest - Monoculture | 0.303 | 0.214 | 0.392 | <0.001 |

**Table S7.** Checklist of avian species detected during the sampling period. We also report the number of individuals of each species detected across land-use types. Species are sorted according to families.

| No. | Family | Species | No.<br>detected<br>in<br>Forests | No.<br>detected in<br>Agroforests | No. detected<br>in<br>Monoculture<br>s |
| --- | --- | --- | --- | --- | --- |
| 1 | Accipitridae | *Tachyspiza badia* | 4 | 3 | 8 |
| 2 | Accipitridae | *Tachyspiza virgata* | 1 | 0 | 0 |
| 3 | Accipitridae | *Haliastur indus* | 0 | 1 | 0 |
| 4 | Accipitridae | *Nisaetus cirrhatus* | 4 | 0 | 0 |
| 5 | Accipitridae | *Pernis ptilorhyncus* | 0 | 1 | 0 |
| 6 | Accipitridae | *Spilornis cheela* | 9 | 4 | 10 |
| 7 | Acrocephalidae | *Acrocephalus*<br>*dumetorum* | 46 | 71 | 118 |
| 8 | Aegithinidae | *Aegithina tiphia* | 12 | 23 | 37 |
| 9 | Alcedinidae | *Ceyx erithaca* | 1 | 0 | 0 |
| 10 | Alcedinidae | *Halcyon*<br>*smyrnensis* | 4 | 9 | 24 |
| 11 | Bucerotidae | *Anthracoceros*<br>*coronatus* | 14 | 14 | 9 |
| 12 | Bucerotidae | *Buceros bicornis* | 3 | 1 | 0 |
| 13 | Bucerotidae | *Ocyceros birostris* | 3 | 0 | 0 |
| 14 | Bucerotidae | *Ocyrceros griseus* | 52 | 18 | 4 |
| 15 | Campephagidae | *Pericrocotus*<br>*cinnamomeus* | 7 | 11 | 9 |
| 16 | Campephagidae | *Pericrocotus*<br>*flammeus* | 0 | 6 | 0 |
| 17 | Charadriidae | *Vanellus indicus* | 0 | 5 | 3 |
| 18 | Chloropseidae | *Chloropsis*<br>*aurifrons* | 2 | 15 | 12 |
| 19 | Chloropseidae | *Chloropsis jerdoni* | 0 | 3 | 0 |
| 20 | Cisticolidae | *Orthotomus*<br>*sutorius* | 12 | 61 | 102 |
| 21 | Cisticolidae | *Prinia hodgsonii* | 1 | 2 | 44 |
| 22 | Cisticolidae | *Prinia inornata* | 0 | 9 | 20 |
| 23 | Cisticolidae | *Prinia socialis* | 0 | 17 | 65 |
| 24 | Columbidae | *Chalcophaps indica* | 52 | 6 | 15 |
| 25 | Columbidae | *Ducula cuprea* | 2 | 0 | 0 |
| 26 | Columbidae | *Spilopelia chinensis* | 0 | 39 | 61 |
| 27 | Columbidae | *Treron affinis* | 62 | 16 | 9 |
| 28 | Corvidae | *Corvus macrorhynchos* | 26 | 35 | 58 |
| 29 | Corvidae | *Corvus splendens* | 0 | 0 | 2 |
| 30 | Corvidae | *Dendrocitta vagabunda* | 1 | 16 | 23 |
| 31 | Cuculidae | *Cacomantis passerinus* | 1 | 3 | 7 |
| 32 | Cuculidae | *Centropus sinensis* | 22 | 56 | 102 |
| 33 | Cuculidae | *Hierococcyx varius* | 9 | 17 | 30 |
| 34 | Cuculidae | *Eudynamys scolopaceus* | 0 | 11 | 18 |
| 35 | Dicaeidae | *Pachyglossa agilis* | 13 | 24 | 45 |
| 36 | Dicaeidae | *Dicaeum concolor* | 57 | 50 | 69 |
| 37 | Dicaeidae | *Dicaeum erythrorhynchus* | 1 | 2 | 4 |
| 38 | Dicruridae | *Dicrurus aeneus* | 29 | 17 | 6 |
| 39 | Dicruridae | *Dicrurus caerulescens* | 0 | 0 | 6 |
| 40 | Dicruridae | *Dicrurus hottentottus* | 6 | 0 | 1 |
| 41 | Dicruridae | *Dicrurus leucophaeus* | 44 | 45 | 67 |
| 42 | Dicruridae | *Dicrurus paradiseus* | 104 | 87 | 121 |
| 43 | Estrildidae | *Lonchura striata* | 0 | 2 | 1 |
| 44 | Fringillidae | *Carpodacus erythrinus* | 1 | 0 | 2 |
| 45 | Irenidae | *Irena puella* | 2 | 0 | 0 |
| 46 | Leiothrichidae | *Alcippe poioicephala* | 88 | 99 | 132 |
| 47 | Leiothrichidae | *Dumetia atriceps* | 13 | 4 | 0 |
| 48 | Leiothrichidae | *Argya striata* | 3 | 20 | 58 |
| 49 | Leiothrichidae | *Argya subrufa* | 0 | 2 | 0 |
| 50 | Megalaimidae | *Psilopogon haemacephalus* | 9 | 8 | 11 |
| 51 | Megalaimidae | *Psilopogon malabaricus* | 3 | 0 | 3 |
| 52 | Megalaimidae | *Psilopogon viridis* | 77 | 115 | 135 |
| 53 | Megalaimidae | *Psilopogon zeylanicus* | 1 | 3 | 4 |
| 54 | Meropidae | *Merops*<br>*leschenaulti* | 0 | 4 | 0 |
| 55 | Meropidae | *Merops orientalis* | 2 | 4 | 21 |
| 56 | Monarchidae | *Hypothymis azurea* | 29 | 11 | 4 |
| 57 | Monarchidae | *Terpsiphone*<br>*paradisi* | 16 | 7 | 3 |
| 58 | Muscicapidae | *Copsychus*<br>*malabaricus* | 20 | 2 | 1 |
| 59 | Muscicapidae | *Copsychus saularis* | 1 | 16 | 17 |
| 60 | Muscicapidae | *Cyornis pallidipes* | 3 | 0 | 0 |
| 61 | Muscicapidae | *Cyornis*<br>*rubeculoides* | 0 | 2 | 0 |
| 62 | Muscicapidae | *Cyornis tickelliae* | 4 | 25 | 18 |
| 63 | Muscicapidae | *Eumyias*<br>*thalassinus* | 0 | 1 | 0 |
| 64 | Muscicapidae | *Muscicapa dauurica* | 1 | 2 | 2 |
| 65 | Muscicapidae | *Muscicapa muttui* | 1 | 1 | 0 |
| 66 | Muscicapidae | *Myophonus horsfieldii* | 23 | 11 | 3 |
| 67 | Muscicapidae | *Saxicola caprata* | 0 | 0 | 1 |
| 68 | Nectariniidae | *Arachnothera longirostra* | 3 | 7 | 1 |
| 69 | Nectariniidae | *Cinnyris asiaticus* | 56 | 105 | 378 |
| 70 | Nectariniidae | *Leptocoma minima* | 196 | 172 | 208 |
| 71 | Nectariniidae | *Leptocoma zeylonica* | 7 | 15 | 14 |
| 72 | Oriolidae | *Oriolus kundoo* | 4 | 13 | 6 |
| 73 | Oriolidae | *Oriolus xanthornus* | 68 | 111 | 187 |
| 74 | Passeridae | *Gymnoris*<br>*xanthocollis* | 0 | 0 | 1 |
| 75 | Pellorneidae | *Pellorneum*<br>*ruficeps* | 18 | 9 | 19 |
| 76 | Phasianidae | *Gallus sonneratii* | 34 | 26 | 62 |
| 77 | Phasianidae | *Pavo cristatus* | 3 | 11 | 29 |
| 78 | Phylloscopidae | *Phylloscopus*<br>*magnirostris* | 17 | 0 | 0 |
| 79 | Phylloscopidae | *Phylloscopus*<br>*nitidus* | 36 | 31 | 25 |
| 80 | Phylloscopidae | *Phylloscopus*<br>*trochiloides* | 9 | 3 | 2 |
| 81 | Picidae | *Micropternus*<br>*brachyurus* | 9 | 6 | 6 |
| 82 | Picidae | *Chrysocolaptes*<br>*socialis* | 13 | 2 | 0 |
| 83 | Picidae | *Yungipicus nanus* | 0 | 0 | 1 |
| 84 | Picidae | *Dinopium benghalense* | 26 | 31 | 19 |
| 85 | Picidae | *Hemicircus canente* | 2 | 1 | 0 |
| 86 | Psittaculidae | *Loriculus vernalis* | 3 | 4 | 2 |
| 87 | Psittaculidae | *Psittacula cyanocephala* | 2 | 7 | 6 |
| 88 | Psittaculidae | *Psittacula krameri* | 1 | 0 | 0 |
| 89 | Pycnonotidae | *Acritillas indica* | 187 | 31 | 26 |
| 90 | Pycnonotidae | *Pycnonotus cafer* | 2 | 32 | 119 |
| 91 | Pycnonotidae | *Pycnonotus jocosus* | 37 | 165 | 399 |
| 92 | Pycnonotidae | *Rubigula gularis* | 0 | 2 | 0 |
| 93 | Pycnonotidae | *Microtarsus priocephalus* | 16 | 11 | 6 |
| 94 | Rhipiduridae | *Rhipidura albogularis* | 1 | 0 | 0 |
| 95 | Sittidae | *Sitta frontalis* | 1 | 0 | 0 |
| 96 | Strigidae | *Glaucidium radiatum* | 9 | 3 | 0 |
| 97 | Timaliidae | *Pomatorhinus horsfieldii* | 39 | 30 | 45 |
| 98 | Trogonidae | *Harpactes fasciatus* | 32 | 0 | 0 |
| 99 | Turdidae | *Turdus simillimus* | 0 | 1 | 1 |
| 100 | Turdidae | *Geokichla citrina* | 3 | 19 | 21 |
| 101 | Vangidae | *Hemipus picatus* | 0 | 1 | 0 |
| 102 | Vangidae | *Tephrodornis pondicerianus* | 0 | 2 | 4 |

**Table S8.** Results of the General Linear Model (Gaussian error distribution) to assess the relative influence of local- (proportion of native trees and canopy cover) and landscape-level (forest cover in 800 m buffer radius, and cashew cover in 800 m buffer radius) predictors on Hill-Shannon diversity (taxonomic diversity) of birds.

| Predictor | Estimate | SE | *t* | *p* |
| --- | --- | --- | --- | --- |
| Intercept | 13.628 | 1.890 | 7.207 | <0.001 |
| Canopy cover | -0.014 | 0.022 | -0.669 | 0.504 |
| Proportion of native trees | 3.150 | 1.227 | 2.567 | 0.0118 |
| Forest cover in 800 m buffer | 0.027 | 0.024 | 1.137 | 0.258 |
| Cashew cover in 800 m buffer | 0.062 | 0.044 | 1.39 | 0.168 |

**Table S9.** Results of the General Linear Model (Gaussian error distribution) to assess the relative influence of local- (proportion of native trees, and canopy cover) and landscape-level (forest cover in 800 m buffer radius, and cashew cover in 800 m buffer radius) predictors on SES-pMPD (phylogenetic diversity).

| Predictor | Estimate | SE | *t* | *p* |
| --- | --- | --- | --- | --- |
| Intercept | 0.296 | 0.661 | 0.448 | <0.01 |
| Canopy cover | -0.001 | 0.008 | -0.222 | 0.824 |
| Proportion of native trees | 0.850 | 0.429 | 1.983 | **0.0503** |
| Forest cover in 800 m buffer | 0.001 | 0.008 | 0.207 | 0.836 |
| Cashew cover in 800 m buffer | -0.024 | 0.016 | -1.530 | 0.129 |

**Table S10.** Results of the General Linear Model (Gaussian error distribution) to assess the relative influence of local- (proportion of native trees, and canopy cover) and landscape-level (forest cover in 800 m buffer radius, and cashew cover in 800 m buffer radius) predictors on SES-fMPD (functional diversity).

| **Predictor** | **Estimate** | **SE** | ***t*** | ***p*** |
| --- | --- | --- | --- | --- |
| Intercept | -0.357 | 0.622 | -0.574 | **<0.001** |
| Canopy cover | 0.002 | 0.007 | 0.370 | 0.712 |
| Proportion of native trees | -0.090 | 0.404 | -0.225 | 0.823 |
| Forest cover in 800 m buffer | 0.006 | 0.008 | 0.763 | 0.447 |
| Cashew cover in 800 m buffer | -0.002 | 0.014 | -0.163 | 0.870 |

**Table S11.**
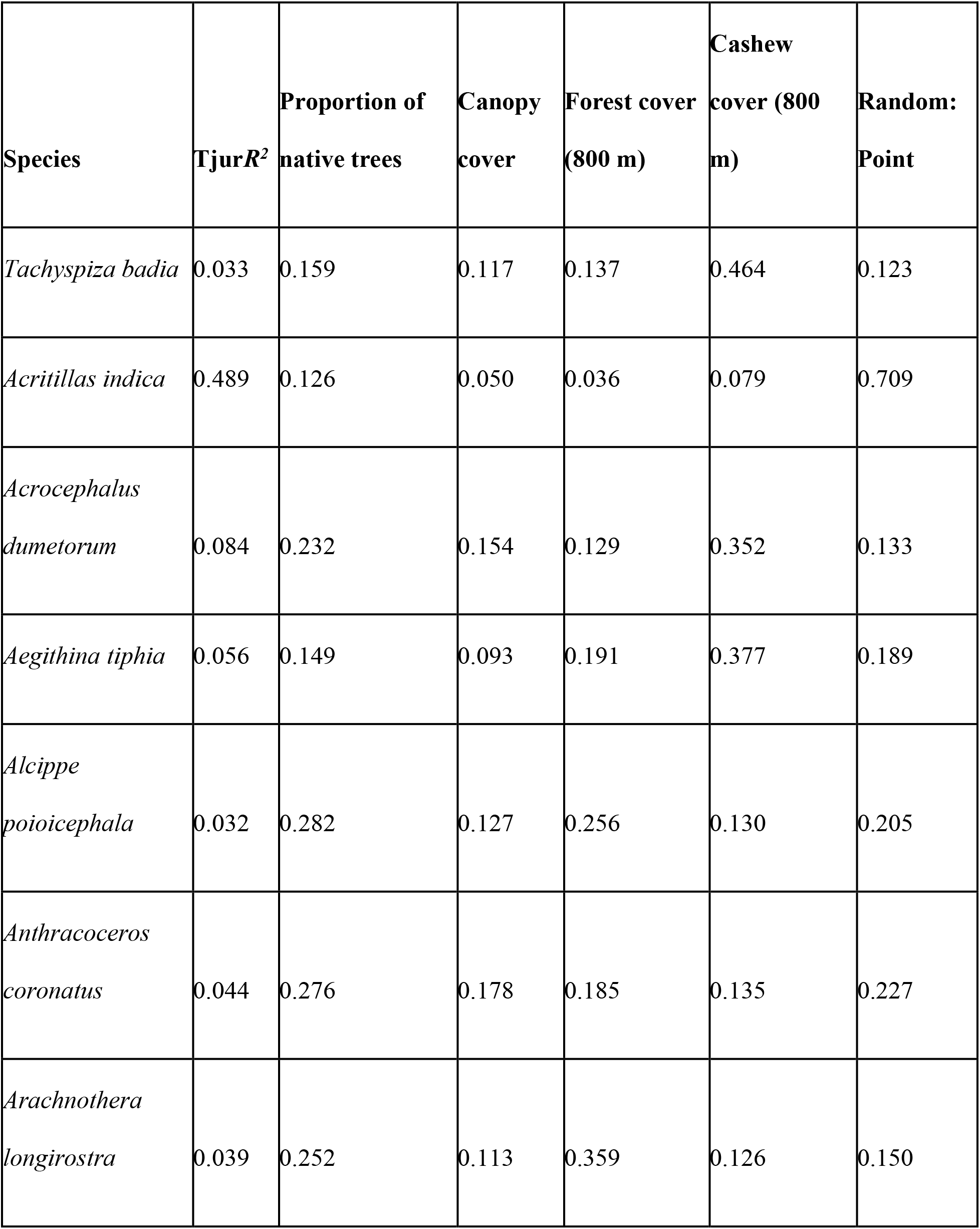
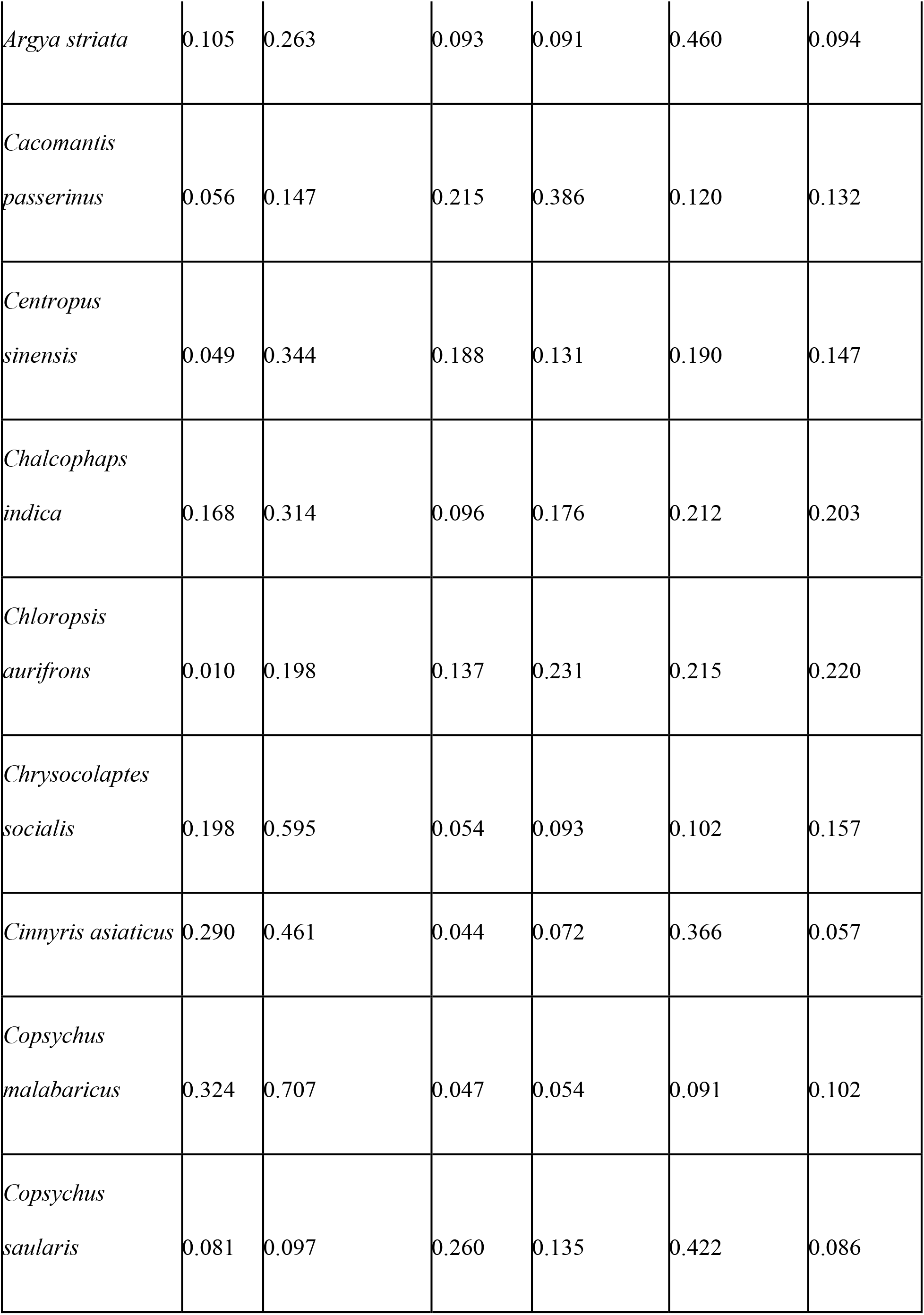
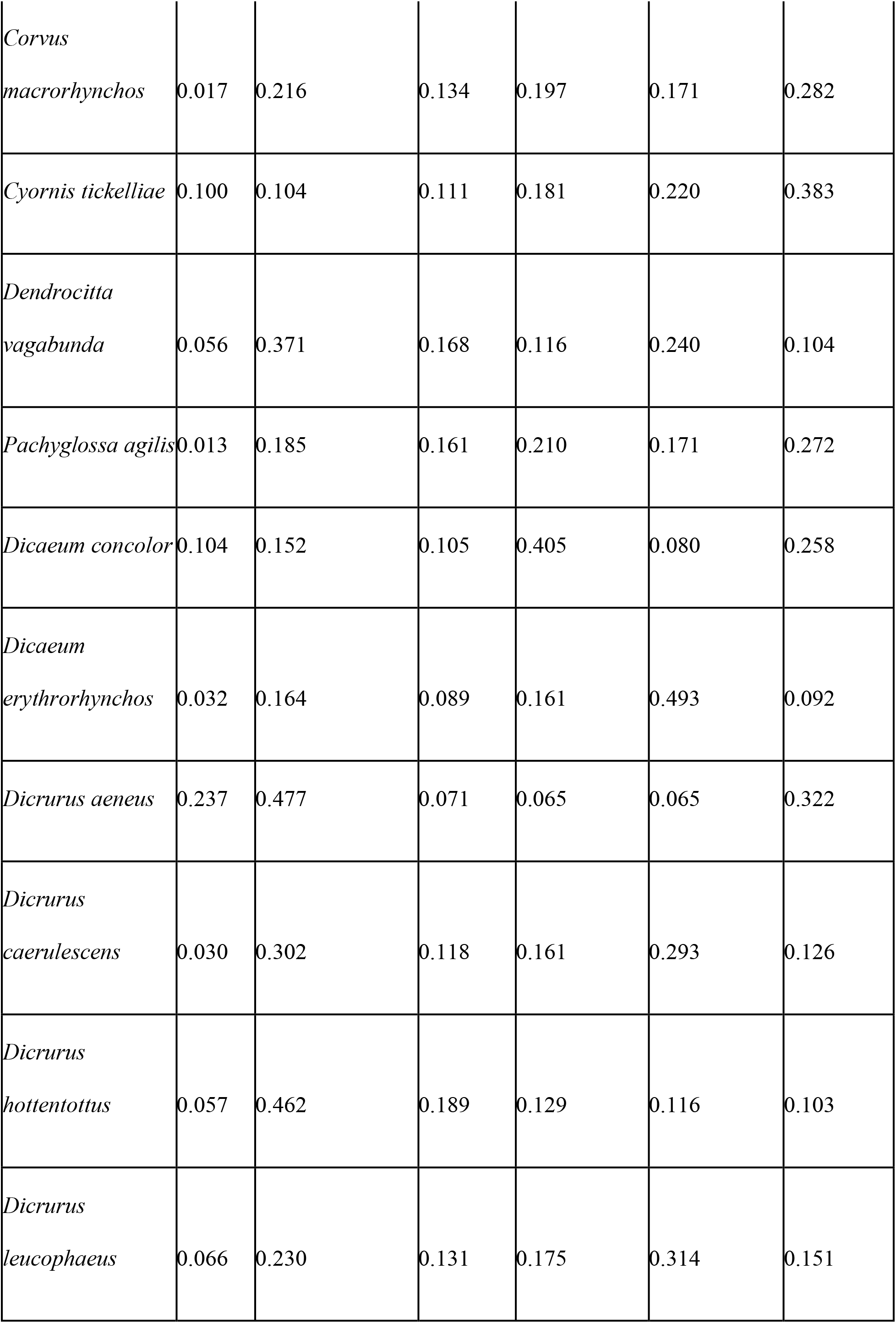
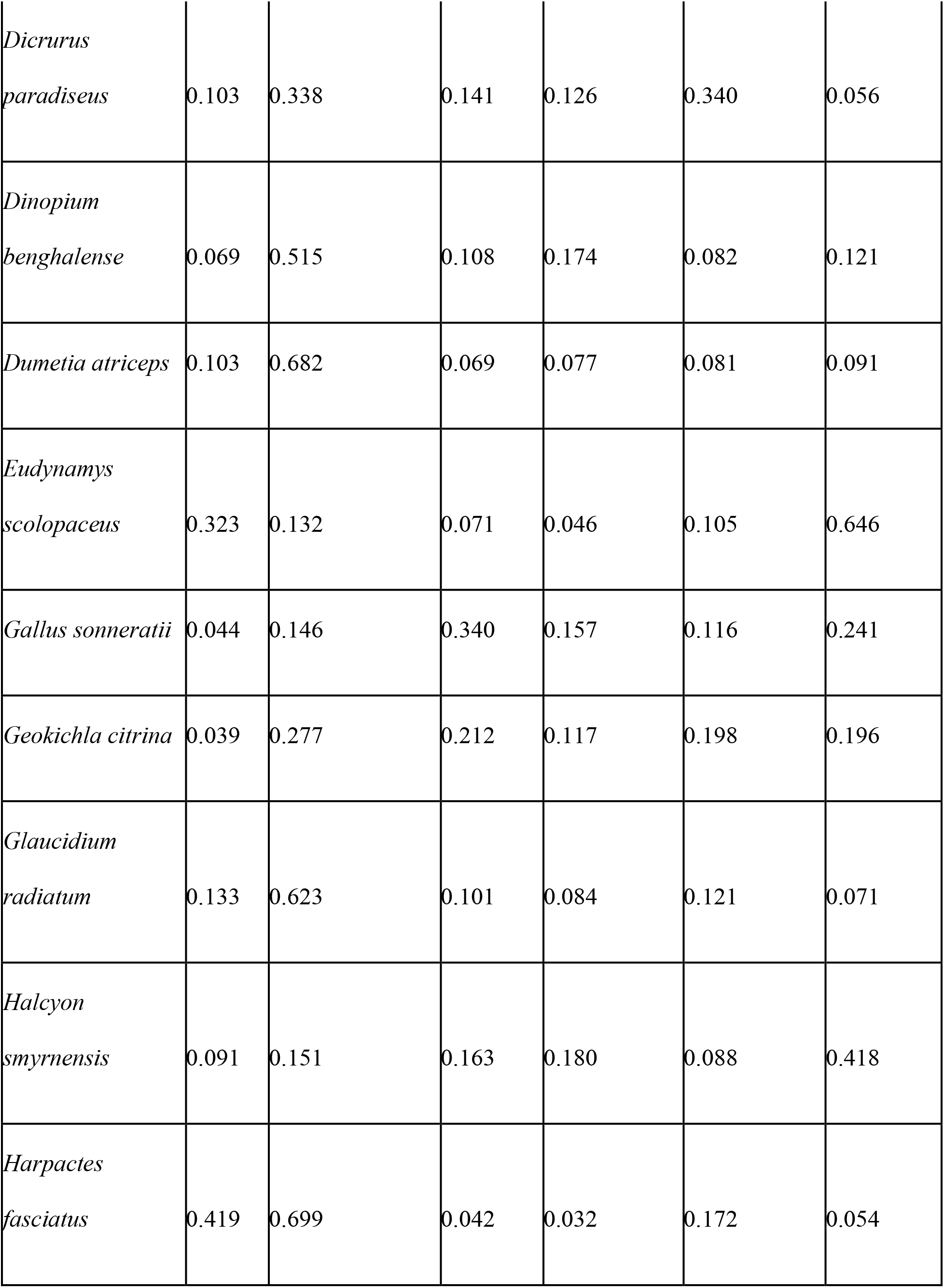
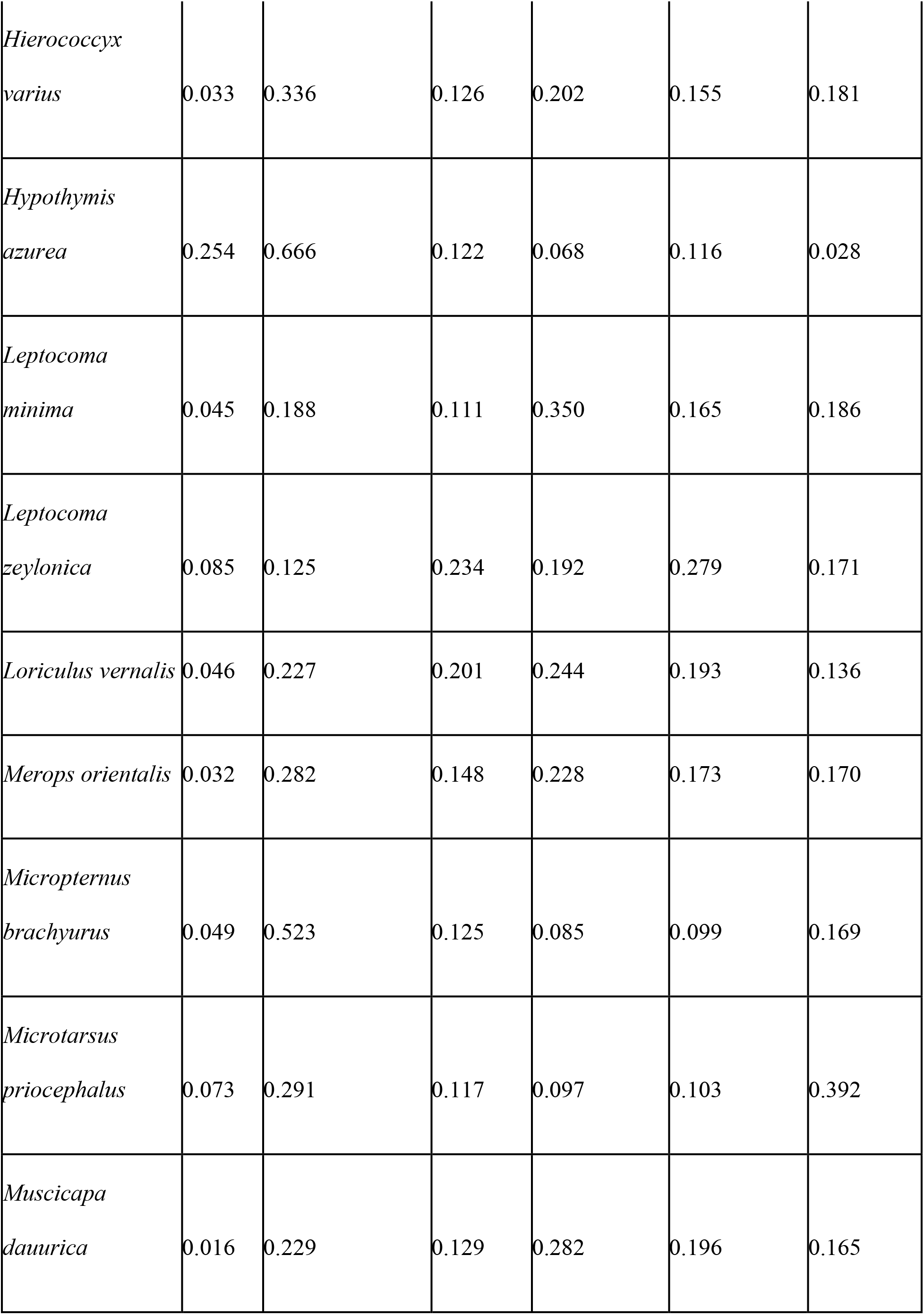
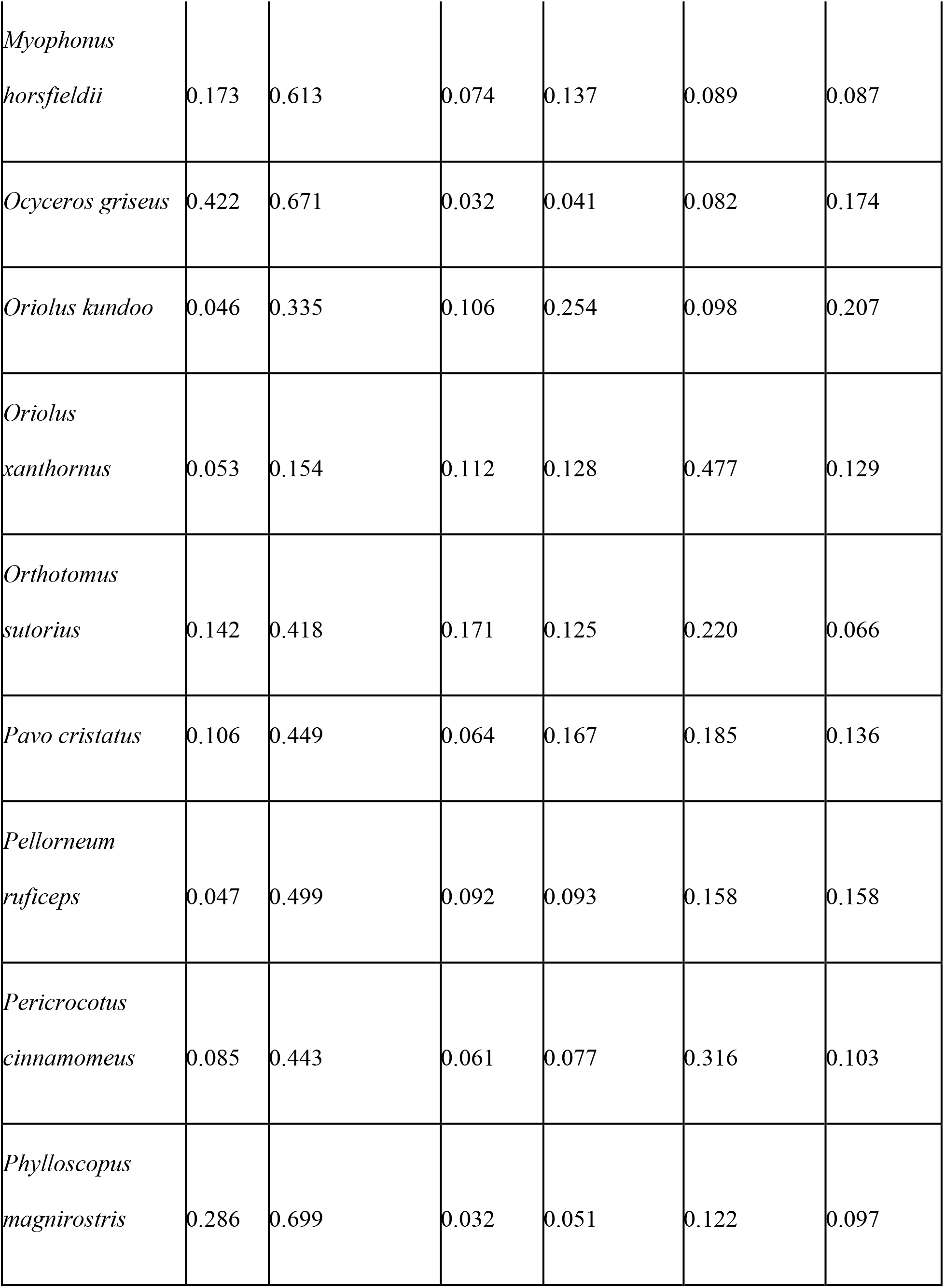
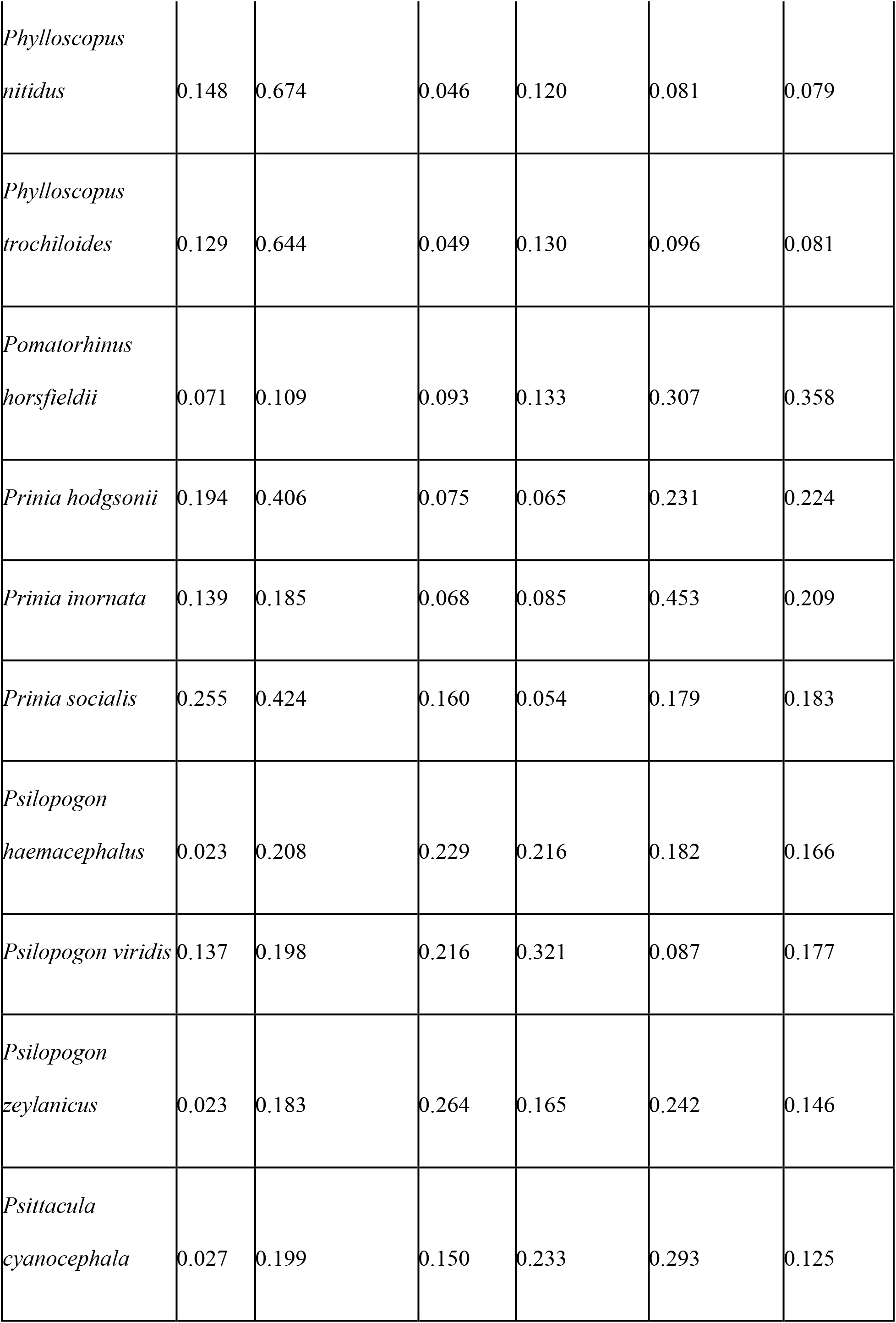
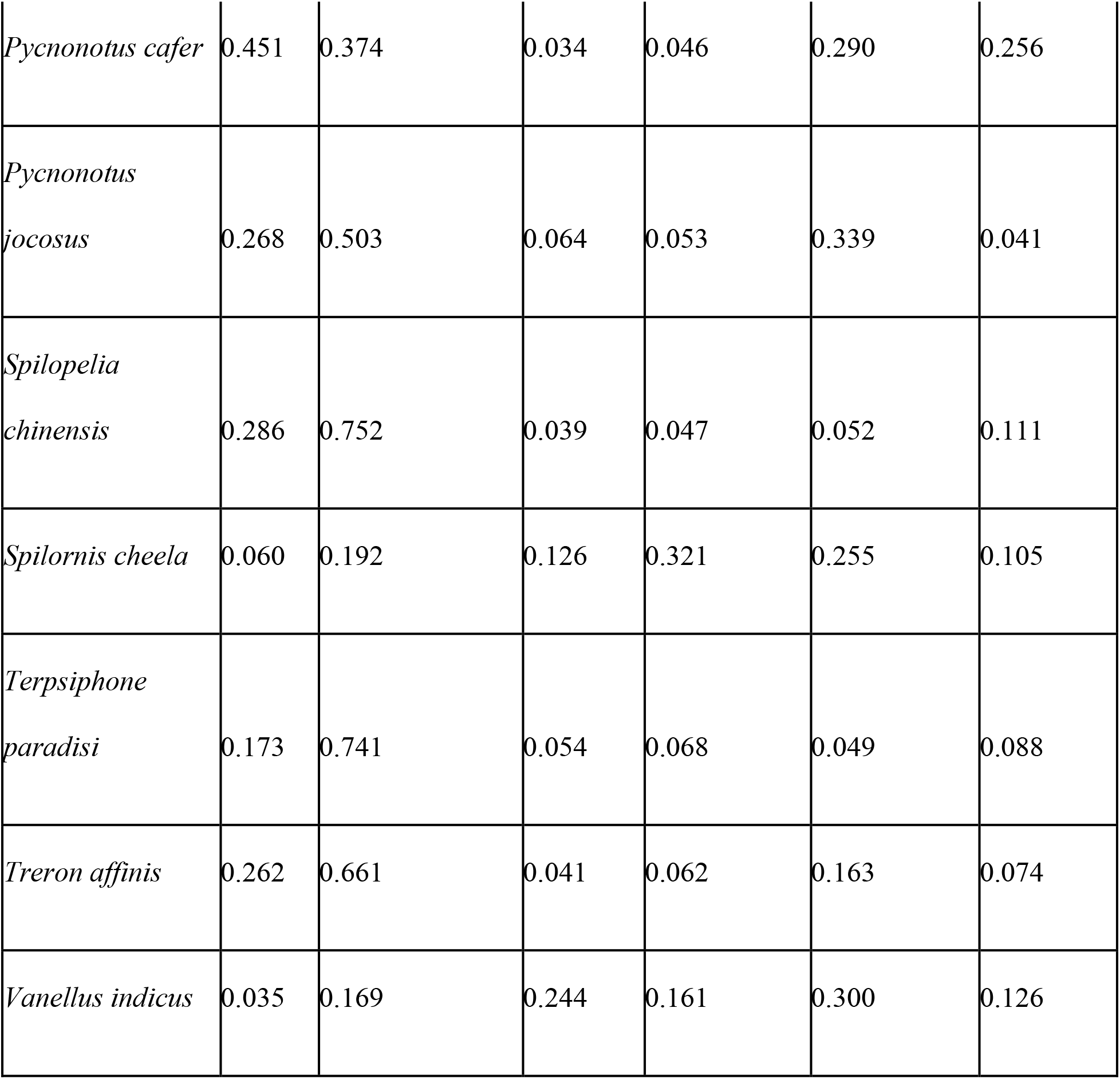
Summary of species-wise Tjur *R^2^* and the proportion of the explained variation by each of the fixed effects (proportion of native trees, canopy cover, forest cover in 800 m and cashew cover in 800 m) and spatial random effect for the 70 species analysed using the Hierarchical Modelling of Species Communities.

